# Inhibition of JAK2V617F-Driven Neoplastic Endothelial Dysfunction by IFNα as a Novel Antifibrotic Mechanism

**DOI:** 10.1101/2024.12.23.630035

**Authors:** Madeline J. Caduc, Mohamed H. E. Mabrouk, Martin Grasshoff, Pia Wanner, Katrin Götz, Jessica E. Pritchard, Dickson W. L. Wong, Cristina B. Lopez, Louisa Böttcher, Pinar Sönmez, Bärbel Junge, Eva M. Buhl, Sabrina Ernst, Gerhard Müller-Newen, Michael Vogt, Jörg Eschweiler, Peter Boor, Maximilian Ackermann, Nicolas Chatain, Julia Franzen, Ivan G. Costa, Martin Zenke, Rebekka K. Schneider, Radek Skoda, Tim H. Brümmendorf, Steffen Koschmieder, Marcelo A. S. de Toledo

## Abstract

The vascular niche is a critical regulator of hematopoiesis and disease progression in myeloproliferative neoplasms (MPN). The presence of JAK2V617F+ endothelial cells (EC) in MPN patients and their association with cardiovascular complications highlight the need to understand and therapeutically target this compartment. Using patient-specific induced pluripotent stem cells (iPSC) harboring JAK2^WT^ or the MPN-driver JAK2V617F (heterozygous, JAK2V617F^HET^, or homozygous, JAK2V617F^HOM^), we identified zygosity-dependent transcriptional profiles in iPSC-derived EC (iEC) at baseline and following interferon-alpha (IFNα) treatment. JAK2V617F^HET^ iEC exhibited an endothelial-to-mesenchymal transition (EndMT) signature, while JAK2V617F^HOM^ iEC showed suppression of translation and ribosome biogenesis. Leveraging iPSC-based 3D assembloids that mimic the bone marrow (BM) niche, we showed that JAK2V617F-driven EndMT is inhibited by tyrosine kinase inhibitors and IFNα. In both JAK2V617F-driven polycythemia vera and TPO-driven myelofibrosis murine models, scRNA-seq analysis of the BM vascular niche consistently revealed inflammatory and EndMT-associated signatures in arterial and arteriolar EC. Notably, dysregulation of ribosome- and translation-related pathways emerged in the myelofibrosis model and at advanced disease stages in JAK2V617F-driven polycythemia vera, indicating progressive vascular remodeling with disease evolution. Chronic pegylated IFNα treatment *in vivo* effectively reversed these pathological changes. IFNα’s anti-EndMT activity was further validated in BM biopsies from MPN patients undergoing IFNα therapy. This is the first study to define MPN stage-dependent vascular remodeling and zygosity-specific endothelial effects of JAK2V617F, and to directly link IFNα-mediated EndMT inhibition as a novel antifibrotic mechanism. Our 3D assembloids provide a translational platform for mechanistic studies and therapeutic targeting of the BM microenvironment in MPN.

**Bullet Points:**

- Arterial vascular remodeling emerges as a novel hallmark of MPN, characterized by TNFα-inflammation, ribosomal dysregulation and EndMT.
- IFNα restores neoplastic endothelial dysfunction, highlighting its role as a vascular niche-modulating and anti-fibrotic agent in MPN.

## Introduction

Myelofibrosis (MF) represents an advanced stage of BCR:ABL1-negative myeloproliferative neoplasms (MPN), marked by progressive bone marrow (BM) fibrosis that impairs normal hematopoiesis and promotes inflammatory and vascular complications^1^. Disruption of the vascular niche, including aberrant neoangiogenesis^2,3^ and endothelial-to-mesenchymal transition (EndMT)^4^, has been observed in MPN progression^5^ and cardiovascular dysfunction (CVD)^6^. While the pathogenesis of cardiovascular complications in MPN remains debated, the detection of JAK2V617F+ endothelial cells (EC) in patients and multilineage analyses suggest that these clonal hematologic malignancies may originate from a common precursor, the hemangioblast^7–10^.

Clinically, in MPN, interferon-alpha (IFNα) induces hematologic and molecular responses and, potentially, delay of fibrotic progression^11,12^. In addition to its antiproliferative and immunomodulatory effects, IFNα also exhibits antitumor properties through inhibition of angiogenesis^13^. While previous studies have described its transcriptional effects on EC^14^, the impact of IFNα on the malignant clone-associated microenvironment, particularly the JAK2V617F+ vascular niche, remains largely unexplored.

Induced pluripotent stem cell (iPSC)-derived organoids and assembloids offer a robust platform for modeling hematologic diseases and drug screening, faithfully recapitulating tissue architecture, cellular composition, and patient-specific genetics *in vitro*^15,16^. In this study, we conducted global transcriptomic profiling of MPN patient-derived iPSC-endothelial cells (iEC) to uncover zygosity-specific transcriptional programs and distinct responses to IFNα treatment. To further dissect the neoplastic vascular niche and its modulation by IFNα, we employed single-cell RNA sequencing (scRNA-seq) of complex iPSC-derived assembloids composed of wild-type or JAK2V617F+ endothelial and hematopoietic cells. These *in vitro* findings were complemented by *in vivo* analyses using two distinct murine MPN models: a JAK2V617F-driven polycythemia vera (PV) model and a thrombopoietin (TPO) overexpression model that recapitulates advanced MF. Both models included treatment with pegylated IFNα (pIFNα) or vehicle to evaluate EndMT and microenvironmental transcriptomic remodeling at single-cell resolution. Finally, the anti-EndMT activity of pIFNα was validated in BM biopsies from MPN patients undergoing pIFNα therapy.

We provide the first evidence of IFNα-mediated regulation of neoplastic endothelial dysfunction in MPN across both human and murine disease models, as well as patient material. Moreover, our 3D *ex vivo* model provides a physiologically relevant platform to dissect cellular crosstalk underlying MF pathogenesis and to evaluate therapeutic effects on the BM niche.

## Methods

### Ethics Approval and Consent to Participate

Peripheral blood mononuclear cells from MPN patients were obtained from the centralized Biomaterial Bank of the Faculty of Medicine, RWTH Aachen University Medical Center. The mesenchymal stromal cells (MSC) used in this study were isolated from the femoral heads of patients undergoing hip replacement surgery (Department of Orthopedics, Trauma and Reconstructive Surgery, Faculty of Medicine, RWTH Aachen University). All biological samples were obtained after written informed consent, as approved by the Ethics Committee of the RWTH Aachen University (EK206/09, EK127/12, EK300/13). Murine experiments were conducted in accordance with institutional guidelines and approved by the relevant ethics committee (81-02.04.2019.A380).

### Differentiation of MPN Patient-Specific iPSC Towards Endothelial and Hematopoietic Lineages

The iPSC lines used in this study were previously characterized^17,18^ and are listed in Supplemental Table 1. Differentiation into hematopoietic and endothelial lineages followed established protocols^19–21^.

### Generation of 3D Fibrin-Based Assembloids

Fibrin hydrogels were generated as previously described^22,23^, with compositions detailed in Supplemental Table 2. Detailed protocols for the 3D experimental setups are provided in the Supplemental Methods.

### MPN Murine Models

The JAK2V617F-driven PV model and TPO-overexpression model of MF have been described previously^24,25^. Detailed methods are provided in the Supplemental Methods.

### Statistical Analysis

Graphical display and statistical analysis were performed with Prism (GraphPad, San Diego, CA, USA). P values of <0.05 were considered statistically significant. Results are presented as mean ± standard deviation (*p < 0.05, **p < 0.01, ***p < 0.001).

### Additional Experimental Procedures

The Supplemental Methods include detailed protocols for iPSC differentiation, generation of iPSC-based assembloids, proliferation and apoptosis assays, *in vitro* drug screening, confocal and transmission electron microscopy, and validation studies using MPN murine models and human BM trephines. Additional protocols for immunofluorescence, flow cytometry, bulk and single-cell RNA sequencing, RT-qPCR, and associated reagents are provided, including IF antibodies (Supplemental Table 3), primers (Supplemental Table 4), FACS antibodies (Supplemental Table 5), compounds (Supplemental Table 6), and media compositions.

## Results

### JAK2V617F Zygosity Shapes Transcriptional Programs in iEC at Baseline and Following IFNα Treatment

To investigate the impact of JAK2V617F on human vasculogenesis, we utilized high-purity (97.13±3.27%) clonal patient-specific JAK2^WT^ and JAK2V617F+ CD31+CD144+CD105+ iEC (Supplemental Fig. 1A). While JAK2V617F did not affect endothelial differentiation, angiogenic potential, or viability in response to drugs used or under investigation for MPN, including IFNα, it conferred a proliferative advantage to iEC (Supplemental Fig. 1B-E). Acute IFNα exposure did not affect capillary-like structure formation, whereas pre-sensitization markedly impaired angiogenesis, with no additional effect upon re-exposure (Supplemental Fig. 1F–J).

In global RNA-seq analysis, unsupervised clustering of differentially expressed genes (DEG) segregated samples by JAK2V617F zygosity (Fig. 1A), while JAK2 and STAT1 gene expression levels remained comparable across genotypes (Supplemental Fig. 2A-B, Supplemental Tables 7, 8).

**Figure 1.**
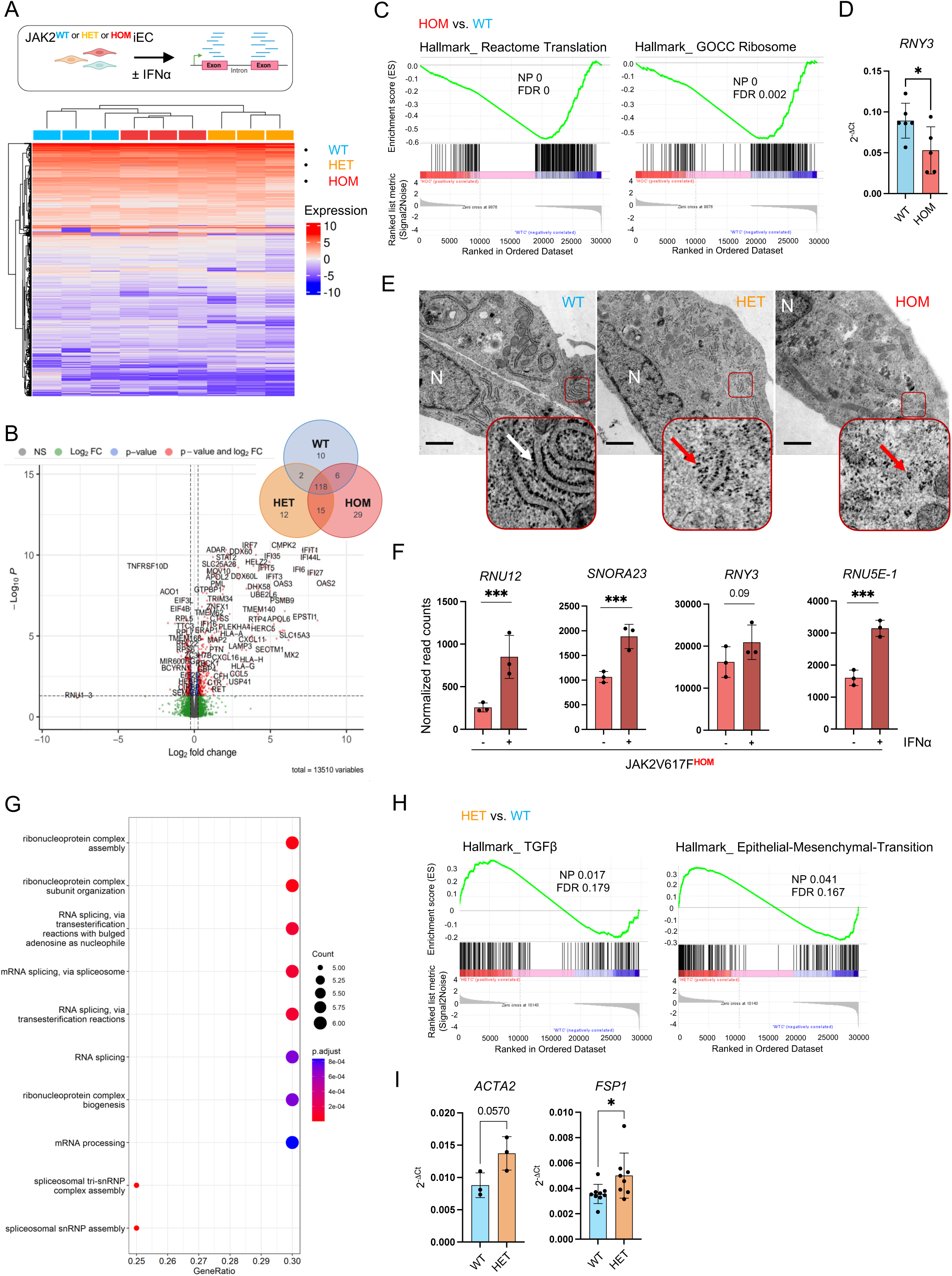
MPN Patient-derived iEC display zygosity-specific transcriptional responses to IFNα treatment (A) Heatmap depicting unsupervised clustering of differentially regulated genes in JAK2^WT^ and JAK2V617F^HET^ ^and^ ^HOM^ iEC (cut off: absolute logarithmic fold change >1). (B) Volcano plot displaying DEG in iEC following IFNα treatment. A total of 168 genes were upregulated, while 10 were downregulated. Abbreviations: NS, non-significant; Log2FC, logarithmic fold change. The accompanying Venn diagram highlights overlapping and significantly enriched genes within each genotype after IFNα treatment (FDR < 0.05). A complete list of all significantly dysregulated genes is provided in Supplemental Table 10. (C) Selection of dysregulated gene sets illustrating the prominent aspects detected in GSEA affecting JAK2V617F^HOM^ iEC biology. Abbreviations: NP, nominal p-value; FDR, false discovery rate. (D) Quantification of mRNA levels of RNA, Ro60-Associated Y3 (*RNY3*) by RT-qPCR in JAK2V617F^HOM^ iEC, relative to JAK2^WT^ iEC. Unpaired t-test (n = 5-6). (E) Representative TEM images of iEC of indicated JAK2V617F zygosity. N marks the nucleus, white arrows depict rough endoplasmic reticulum, and red arrows showcase accumulated loose ribosomes in mutated iEC. Scale bar: 1,000 nm. (F) Quantification of normalized mRNA read counts for RNA, U12 small nuclear (*RNU12*), Small Nucleolar RNA, H/ACA Box 23 (*SNORA23*), *RNY3*, and RNA, U5E small nuclear 1 (*RNU5E-1*) expression in untreated versus IFNα-treated JAK2V617F^HOM^ iEC (n = 3). (G) GO analysis depicting upregulated terms in JAK2V617F^HOM^ iEC following IFNα treatment. The size of the dots reflects the number of dysregulated genes from the gene list associated with the GO term, while the color of the dots represents the adjusted p-value (p. adjust). (H) Selected dysregulated gene sets from GSEA highlighting key pathways altered in JAK2V617F^HET^ iEC. Abbreviations: NP, nominal p-value; FDR, false discovery rate. (I) Quantification of mRNA levels of fibroblast-specific protein 1 (*FSP1*), alpha-smooth muscle actin (*ACTA2*) by RT-qPCR in JAK2V617F^HET^ iEC, relative to JAK2^WT^ iEC. Unpaired t-test (n = 3-8).

Upon IFNα treatment, iEC showed a robust interferon response, with upregulation of antiviral and immunomodulatory pathways (Fig. 1B; Supplemental Fig. 2C). Key interferon-stimulated genes (*IFIT1*, *STAT1*, *USP18*) were validated by RT-qPCR (Supplemental Fig. 2D). IFNα treatment did not induce apoptosis (Supplemental Fig. 2E–F), consistent with drug screening results showing preserved iEC viability after 72 hours treatment (Supplemental Fig. 1E). PROGENy analysis^26^ confirmed similar JAK–STAT activation across genotypes (Supplemental Fig. 2G).

Differential expression analysis revealed zygosity-dependent transcriptional alterations both at baseline and following IFNα treatment. At baseline, JAK2V617F^HOM^ iEC exhibited downregulation of translation- and ribosome-associated gene sets (Fig. 1C), including ribosomal regulatory non-coding RNA (ncRNA) such as SNORA73B (Fig. 1D; Supplemental Fig. 3A)^27^. Disruption of pathways related to ribosome biogenesis, ribonucleoprotein complex assembly, and RNA splicing was also observed (Supplemental Fig. 3B). Consistently, transmission electron microscopy (TEM) revealed disorganized ER–ribosome architecture and reduced rough ER (rER) in homozygous iEC (Fig. 1E; Supplemental Fig. 3C; Supplemental Table 9). Following IFNα treatment, JAK2V617F^HOM^ iEC displayed the highest number of uniquely regulated DEG (Supplemental Table 10), including restoration of ncRNA (Fig. 1F)^28,29^, and enrichment of ribosome biogenesis and mRNA splicing pathways (Fig. 1G). This effect was not observed in JAK2^WT^ controls (Supplemental Fig. 3D).

In JAK2V617F^HET^ iEC, gene set enrichment analysis (GSEA) identified upregulation of TGFβ signaling and EMT-associated genes (Fig. 1H), along with enrichment of hypoxia and reactive oxygen species (ROS)-related genes (Supplemental Fig. 3E), both shown to be critical for MPN pathogenesis^30,31^. Pathways involved in mesenchymal development, and extracellular matrix (ECM) synthesis were also upregulated (Supplemental Fig. 3F). Concordantly, expression of key EndMT markers such as fibroblast-specific protein 1 (FSP1) and α-smooth muscle actin (ACTA2) was increased in JAK2V617F^HET^ ^vs. WT^ iEC (Fig. 1I; Supplemental Fig. 3G-H)^32^. These findings demonstrate that JAK2V617F zygosity drives distinct transcriptional programs in iEC which can be reversed by IFNα treatment.

### IFNα and Tyrosine Kinase Inhibitors Mitigate the Pro-Fibrotic Phenotype of JAK2V617F^HET^ iEC

Tyrosine kinase inhibitors like ruxolitinib and nintedanib exert anti-fibrotic effects, including EndMT suppression^33–35^, yet the mechanisms underlying IFNα’s disease-modifying properties remain unclear^36^. Given the pro-mesenchymal signature observed in JAK2V617F^HET^ iEC, we investigated their contribution in BM niche fibrosis and IFNα’s capacity to counteract EndMT. Ruxolitinib served as a positive control, while nintedanib, an emerging agent in hematologic disorders^37^, was tested alone and in combination with ruxolitinib to assess potential synergy.

We developed 3D fibrin-based assembloids that mimic critical EC-MSC interactions required for vessel formation^38^ (Fig. 2A). iEC formed vessel-like structures, enhanced by MSC, and the model recapitulated our initial endothelial tube formation assay findings, including the anti-angiogenic impact of IFNα pre-sensitization (Supplemental Fig. 4A–C).

**Figure 2.**
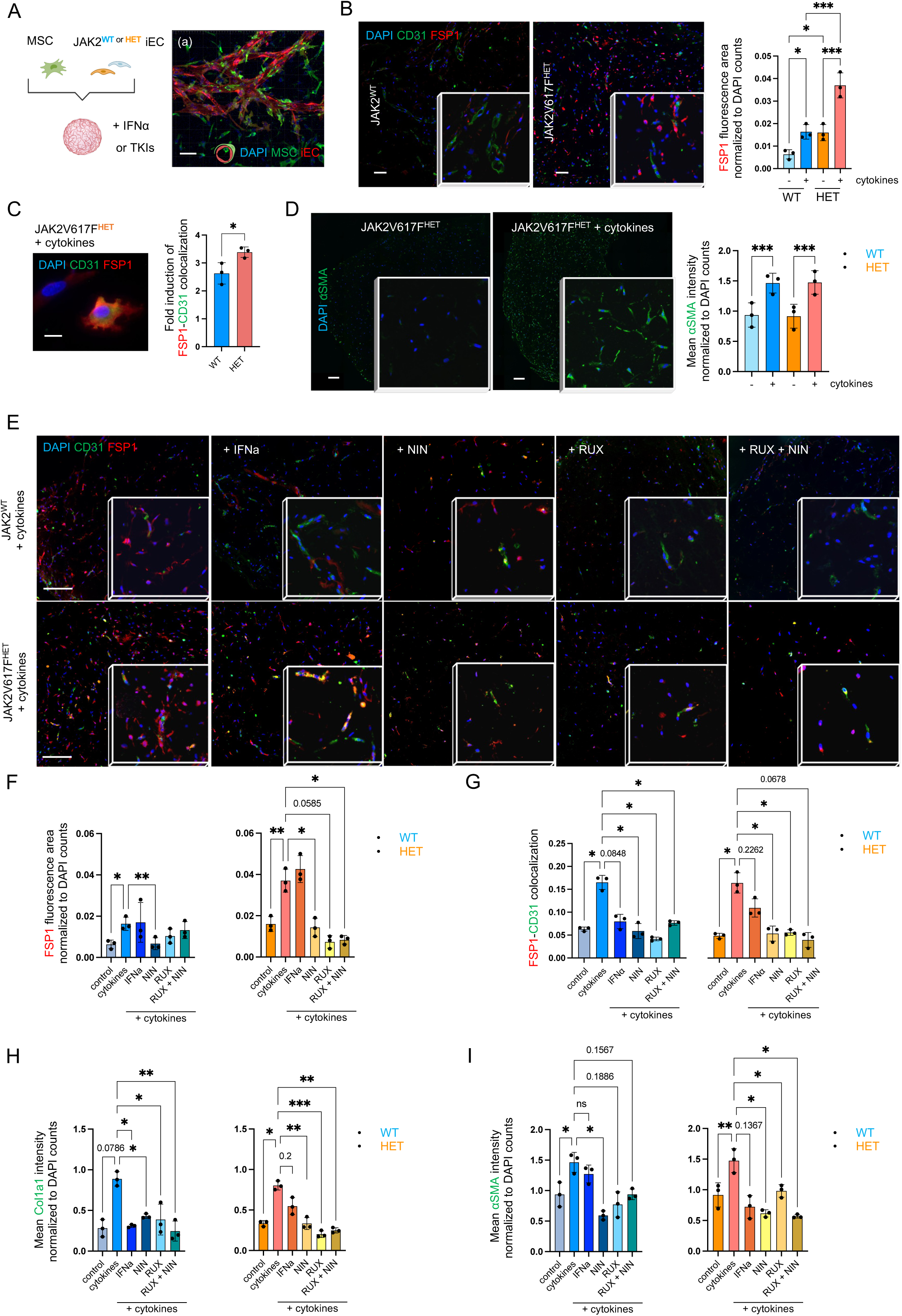
Modeling JAK2V617F-driven EndMT in iPSC-based assembloids (A) Schematic representation of the experimental setup. After vascular tree formation, WT and JAK2V617F^HET^ iEC assembloids were pre-treated with IFNα (1000 U/ml) or with a tyrosine kinase inhibitor (TKI): ruxolitinib (RUX, 250 nM), nintedanib (NIN, 250 nM), or a combination of ruxolitinib and nintedanib (both 250 nM). EndMT was then induced using pro-fibrotic cytokines (IL1β, TNFα, and TGFβ) to mimic the pathological conditions of MF. (a) Representative confocal image shows MSC alignment along the walls of tube-like structures, mimicking the physiological EC-pericyte arrangement. iEC (red) and MSC (green) were labeled with fluorescent cell linkers PKH26 and PKH67, respectively. Nuclei are stained with DAPI. Scale bar: 100 µm. (B) Representative IF of JAK2^WT^ and JAK2V617F^HET^ 3D assembloids, stained for CD31 (green) and FSP1 (red) to indicate EndMT. Cell nuclei are visualized using DAPI (blue). Scale bar: 125 µm. Quantification of FSP1 fluorescence area in JAK2^WT^ and JAK2V617F^HET^ assembloids under both stimulated and unstimulated conditions. Two-way ANOVA (n = 3). (C) Representative duplex IF showing FSP1 expression in a CD31⁺ iEC following exposure to IL1β, TNFα, and TGFβ in JAK2V617F^HET^ 3D assembloids. Scale bar: 10 µm. Fold induction quantification of FSP1-CD31 fluorescence colocalization in JAK2^WT^ vs. JAK2V617F^HET^ 3D assembloids after cytokine stimulation. Unpaired t-test (n = 3). (D) Representative IF of JAK2V617F^HET^ assembloids under basal conditions and after pro-fibrotic cytokine exposure, stained for αSMA (green), with DAPI (blue) highlighting cell nuclei. Quantitative analysis of mean fluorescence intensity (MFI) for αSMA was conducted in mutated and unmutated assembloids. Two-way ANOVA (n = 3). Scale bars: 250 µm. (E) Representative immunofluorescence images of assembloids stained for CD31 (green), FSP1 (red), and nuclei (DAPI, blue) under different treatment conditions. Scale bar: 125 µm. (F) Quantification of FSP1 fluorescence area in cytokine-treated 3D assembloids following sensitization with IFNα, nintedanib, ruxolitinib, or their combination. Statistical analysis by one-way ANOVA (n = 3). (G) Quantification of FSP1–CD31 colocalization in JAK2^WT^ and JAK2V617F^HET^ 3D assembloids under different treatment conditions, using Mander’s colocalization coefficient (M1). One-way ANOVA (n = 3). MFI quantification of pro-fibrotic ECM components (H) Col1a1 and (I) αSMA in 3D assembloids pre-treated with IFNα, nintedanib, ruxolitinib, or their combination, followed by cytokine exposure. One-way ANOVA (n = 3).

Consistent with the pro-mesenchymal transcriptional signature, JAK2V617F^HET^ cocultures showed elevated FSP1 protein levels, both overall and iEC-specific, compared to WT (Fig. 2B–C). Exposure of assembloids to pro-inflammatory and pro-fibrotic cytokines (TGFβ, IL-1β, and TNFα) induced features of myelofibrotic transformation, including increased collagen I (Col1a1) and αSMA deposition (Fig. 2D, Supplemental Fig. 4D). A mesenchymal shift in iEC was confirmed in 2D cultures, with JAK2V617F^HET^ iEC exhibiting reduced *CLDN5* expression and cytokine-induced upregulation of *SNAI1* alongside downregulation of *CD31* (Supplemental Fig. 4E).

Nintedanib, ruxolitinib, and their combination significantly reduced overall and iEC-specific FSP1 protein deposition in JAK2^WT^ and JAK2V617F^HET^ assembloids (Fig. 2E–G), while IFNα induced a trend toward reduction (Fig. 2G). This reduction was associated with decreased Col1a1 and αSMA deposition, while CD31 expression remained unchanged across all conditions, indicating preserved vascular integrity (Fig. 2H–I, Supplemental Fig. 4F). These findings suggest that IFNα and, ruxolitinib and nintedanib, have the potential to suppress EndMT.

### iPSC-Based Assembloids Mimic Key Cellular Interactions of the Human BM Niche

To better replicate *in vivo* conditions and further investigate the role of mutant endothelium and impact of IFNα treatment on BM niche remodeling, we incorporated hematopoietic cells (JAK2^WT^ or JAK2V617F^HET^ iHC) into our assembloids. These included CD117⁺CD34⁺ hematopoietic stem/progenitor cells, CD45⁺CD61/41⁺megakaryocytes, CD45⁺CD68⁺ monocytes/macrophages, and CD45⁺CD66b⁺ granulocytes (Fig. 3A, Supplemental Fig. 5A). scRNAseq of these complex 3D assembloids identified distinct hematopoietic populations based on transcriptional profiles and lineage-defining marker expression (Fig. 3B). Within the non-hematopoietic (CD45⁻) compartment, eight distinct clusters were identified: six mesenchymal, one endothelial, and one hemogenic endothelial cluster (Supplemental Fig. 5B, Supplemental Table 11).

**Figure 3.**
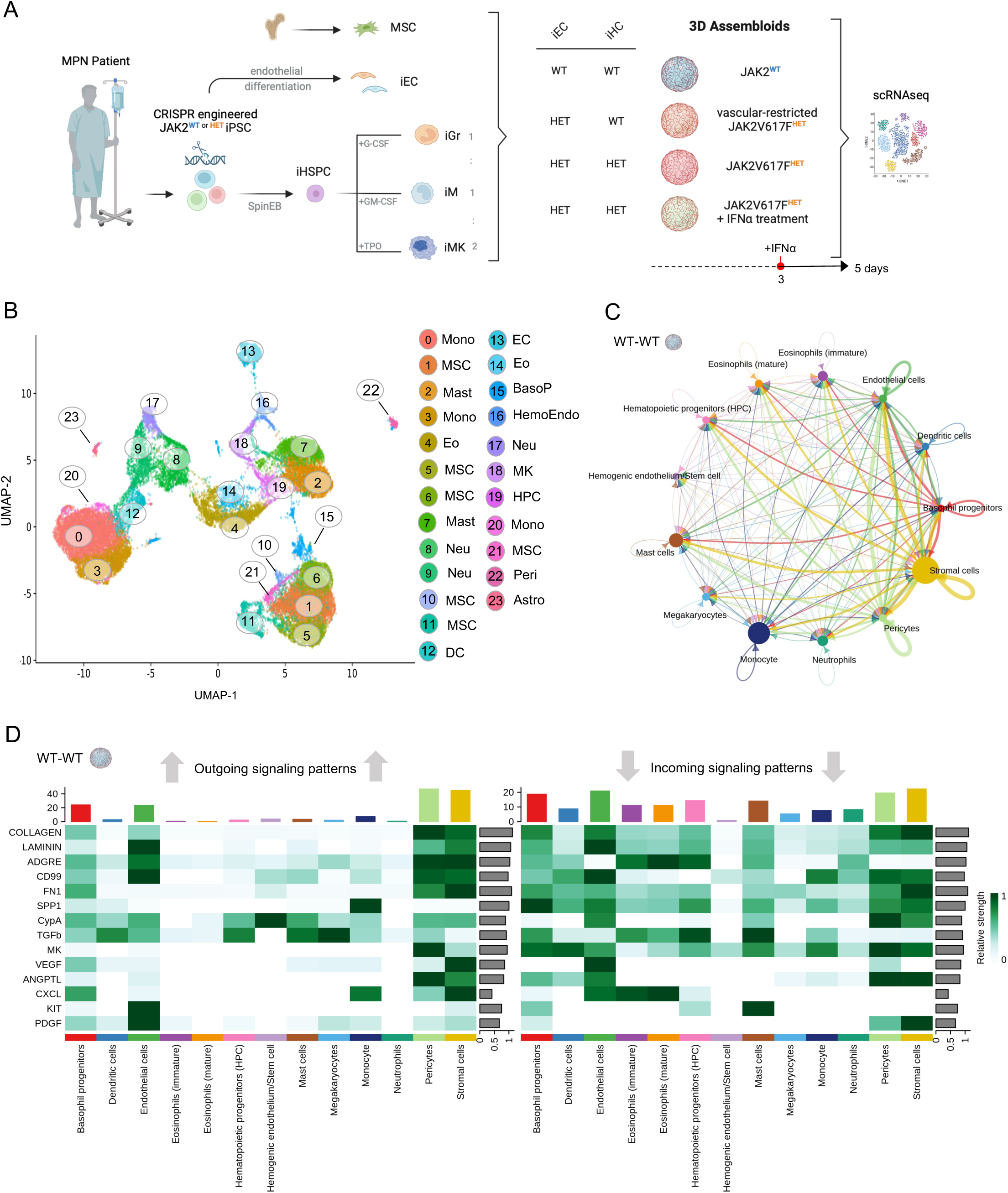
Assembloids mimic key cellular interactions of the human BM niche (A) Schematic of the experimental setup for the generation of 3D assembloids for scRNAseq. JAK2^WT^ or JAK2V617F^HET^ iPSC-derived megakaryocytes (iMK), monocytes/macrophages (iM), and granulocytes (iGr) (collectively referred to as iPSC-derived hematopoietic cells (iHC)) were mixed in a 2:1:1 ratio and combined with MSC and JAK2^WT^ or JAK2V617F^HET^ iEC within fibrin hydrogels. To evaluate the effect of IFNα on the mutant niche, JAK2V617F^HET^ assembloids were treated with IFNα (1000 U/ml) for 48h before scRNAseq. (B) Uniform Manifold Approximation and Projection (UMAP) plot showing the isolated hematopoietic and stromal clusters. Abbreviations: Mono – monocytes; MSC – mesenchymal stromal cells; Mast – mast cells; Eo – eosinophils; Neu – neutrophils; DC – dendritic cells; EC – endothelial cells; BasoP – basophil progenitors; HemoEndo – hemogenic endothelium; MK – megakaryocytes; HPC – hematopoietic progenitor cells; Peri – pericytes; Astro – astrocytes. (C) Circle plot depicting the mapping of ligand-receptor (L-R) interactions among various cell types in JAK2^WT^ (WT-WT) assembloids. Each node represents a distinct cell type, with MSC clusters categorized as stromal cells. Line thickness indicates interaction strength, while arrows denote the direction of the interactions. (D) Heatmap illustrating the relative strength of incoming and outgoing signaling among various cell types in JAK2^WT^ assembloids. Rows represent aggregate strength of signaling for the given pathway, while columns indicate distinct cell types. The color code reflects the strength of the interactions.

We first analyzed scRNA-seq data from unmutated 3D assembloids to map functional crosstalk between cellular subtypes, revealing extensive autocrine and paracrine signaling between stromal and hematopoietic compartments (Fig. 3C). Stromal cells, including EC, and basophil progenitors emerged as primary sources of outgoing signals, engaging in ECM- and growth factor–mediated interactions via ligands such as collagen, laminin, VEGF, PDGF, and KIT (Fig. 3D). Predicted ligand– receptor (L–R) interactions included *ANGPT1/2–TEK*, *KIT–KITL*, *CXCL12–CXCR4*, and *JAG1–NOTCH1*, underscoring the stromal compartment’s central role in supporting BM niche structure and function (Supplemental Fig. 5D).

Together, these findings highlight the transcriptional homology of our assembloids with the human BM niche.

### JAK2V617F Drives Neoplastic Angiogenesis and Dysregulates Cellular Crosstalk in 3D Assembloids

Next, we examined JAK2V617F-driven transcriptional changes, focusing on its effects in EC alone versus in combination with hematopoietic expression. Consistent with our 2D data (Supplemental Fig. 1B), assembloids with JAK2V617F^HET^ endothelium showed a significant increase in EC numbers, which was further amplified by the presence of JAK2V617F^HET^ iHC (Fig. 4A). Mutated iEC exhibited a pro-mesenchymal transcriptional profile, with increased expression of EndMT markers such as *ACTA2*, *FSP1 (S100A4)*, and *FAP* (Fig. 4B), recapitulating our bulk RNA-seq findings (Supplemental Fig. 6A). In assembloids harboring both mutated endothelium and hematopoietic cells, *DEPP1*, *ACTA2*, and *SNAI1* were further upregulated, along with glycolysis-associated genes (*MYC*, *LDH*, *GAPDH*), indicating advanced EndMT (Fig. 4B; Supplemental Fig. 6B)^39^.

**Figure 4.**
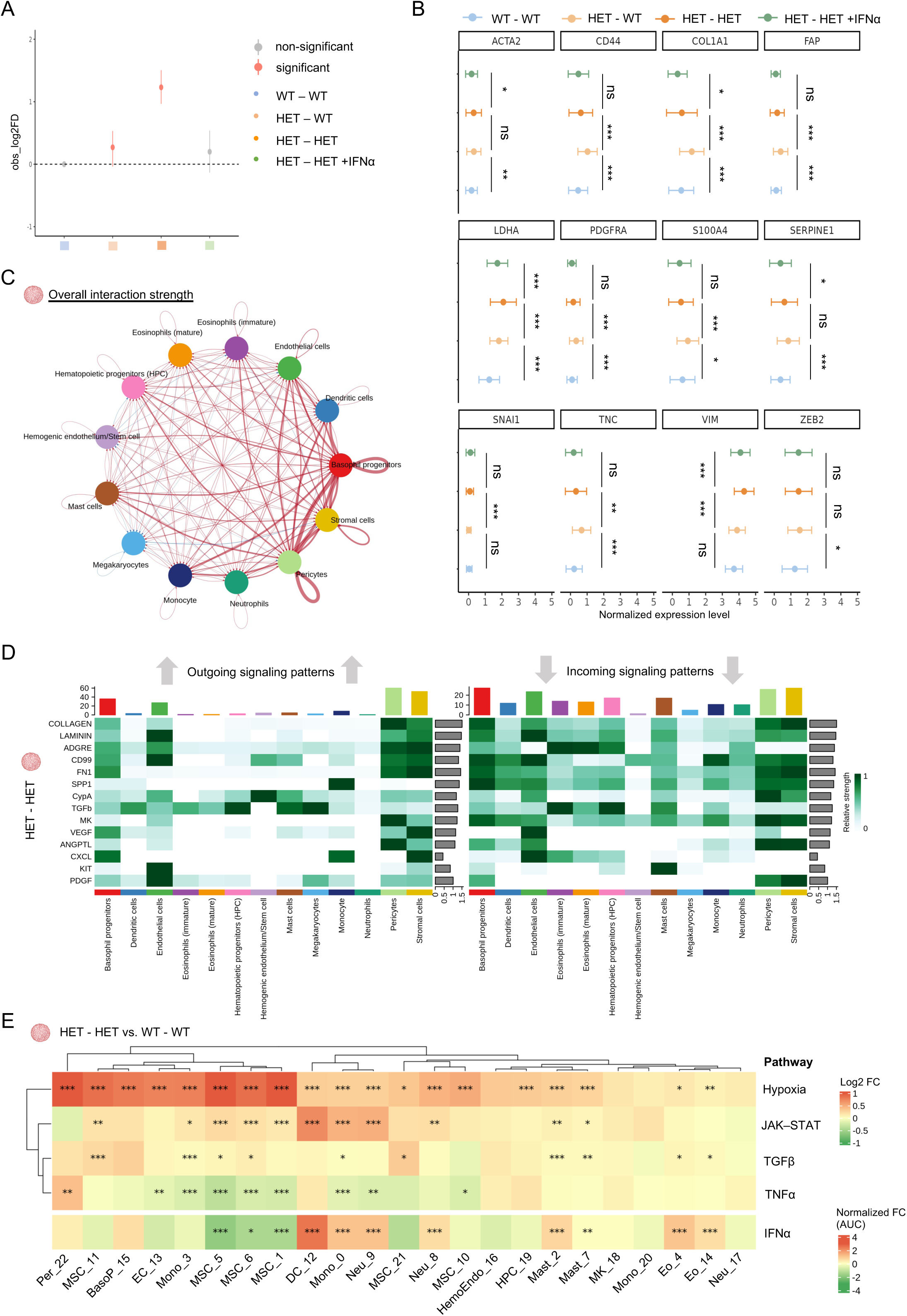
JAK2V617F^HET^ assembloids recapitulate key MPN features (A) Quantification of relative EC numbers in 3D assembloids. (B) Dot plots with error bars showing the normalized expression levels of selected EndMT markers in EC from 3D assembloids. In figure (A) and (B) the color code in the figure represents different genetic backgrounds and treatment conditions: blue for JAK2^WT^ assembloids, yellow for assembloids with vascular-specific JAK2V617F^HET^ expression, orange for JAK2V617F^HET^ assembloids, and green for JAK2V617F^HET^ assembloids treated with IFNα (1000 U/ml) for 48 hours prior to scRNA-seq. (C) Circle plot illustrating the overall ligand-receptor (L-R) interaction strength among various cell types in JAK2V617F^HET^ versus JAK2^WT^ assembloids. Each node represents a distinct cell type, with MSC clusters classified as stromal cells. Line thickness reflects interaction strength, and arrows indicate the direction of interactions. Red lines indicate increased interaction strength, while blue lines denote decreased interaction strength. (D) Heatmap illustrating the relative strength of incoming and outgoing signaling among various cell types in JAK2V617F^HET^ assembloids. Rows represent aggregate strength of signaling for the given pathway, while columns indicate distinct cell types. The color code reflects the strength of the interactions. (E) PROGENy analysis and Area Under the Curve (AUC) score analysis of selected pathways in JAK2V617F^HET^ (HET-HET) compared to JAK2^WT^ assembloids (WT-WT). Abbreviations: FC – Fold change of HET-HET vs. WT-WT assembloids.

While vascular-restricted JAK2V617F^HET^ reduced the strength of EC interactions with niche components (Supplemental Fig. 6C), particularly via *COL4A1/2* and *CALM1*, basophil progenitors exhibited enhanced FN1 signaling through *ITGAV* and *CD44* and stronger interactions with neutrophils, mast cells, and stromal cells via the *SPP1–CD44* axis. Elevated SPP1, CD44, and FN1 are associated with poor cancer prognosis due to their role in tumor microenvironment remodeling^40,41^.

In JAK2V617F^HET^ ^vs. WT^ assembloids, L–R interaction mapping revealed globally increased cellular crosstalk, driven primarily by basophil progenitors and stromal compartments including pericytes: aberrant signaling involved ECM components (e.g., collagen, FN1), angiogenic factors (e.g., PDGF, VEGF, ANGPTL), and inflammatory mediators such as TGFβ (Fig. 4C–D). Additionally, increased interactions were identified between hematopoietic progenitors, megakaryocytes, and EC with basophil progenitors via the *MIF-CD74* axis (Supplemental Fig. 6D), a pathway implicated in MPN development^42^. Moreover, basophil progenitors showed enhanced autocrine and paracrine signaling with megakaryocytes and monocytes via the *LGALS1–ITGB1* axis (Supplemental Fig. 6E), a pro-inflammatory pathway implicated in MF^43^.

PROGENy analysis showed that the presence of JAK2V617F in both endothelial and hematopoietic lineages markedly upregulated hypoxic signaling (Fig. 4E), a hallmark of MPN^31^. Cell types involved in fibrotic remodeling, including monocytes/macrophages, neutrophils, MSC, and mast cells^37,44,45^, exhibited robust activation of the pro-inflammatory JAK-STAT pathway. The TGFβ pathway, a key driver of fibrosis^46^, was activated in MSC populations, mast cells, and monocytes (Fig. 4E). Notably, IFNα and TNFα signaling were suppressed in MSC clusters (1, 5, and 6), whereas hematopoietic cells showed enriched IFNα signaling (Fig. 4E).

Given the key role of the mesenchymal compartment in MF^44,47^, we focused our analysis on the JAK2V617F-driven transcriptional changes in the CD45^-^ populations: MAPK signaling, associated with fibrosis and carcinogenesis^48^, was upregulated in EC (cluster 4) and proliferative MSC (cluster 6), while pericytes (cluster 7) showed enhanced PI3K activation. TGFβ signaling and ECM-related gene expression were selectively enriched in MSC from mutated assembloids, whereas vascular-restricted JAK2V617F predominantly induced ECM remodeling in MSC cluster 3 (Supplemental Fig. 7A–C).

Thus, our assembloids replicate key features of MPN such as hypoxia, inflammation and fibrotic transformation. Moreover, they confirm and extend previous reports^37,49,50^ that basophil progenitors and related cells (e.g., mast cells) may play an important role in MPN pathogenesis.

### IFNα inhibits EndMT progression in JAK2V617F 3D Assembloids

To assess IFNα’s impact on neoplastic vasculature and niche remodeling, JAK2V617F+ assembloids were treated with IFNα for 48h prior to scRNA-seq (Fig. 3A). Consistent with our previous findings, IFNα reversed the hyperproliferative phenotype of mutant iEC (Fig. 4A), downregulated EndMT-associated markers (*ACTA2*, *COL1A1*, *Vimentin*) (Fig. 4B) and reduced endothelial outgoing signaling via laminin and VEGF (Fig. 5A).

**Figure 5.**
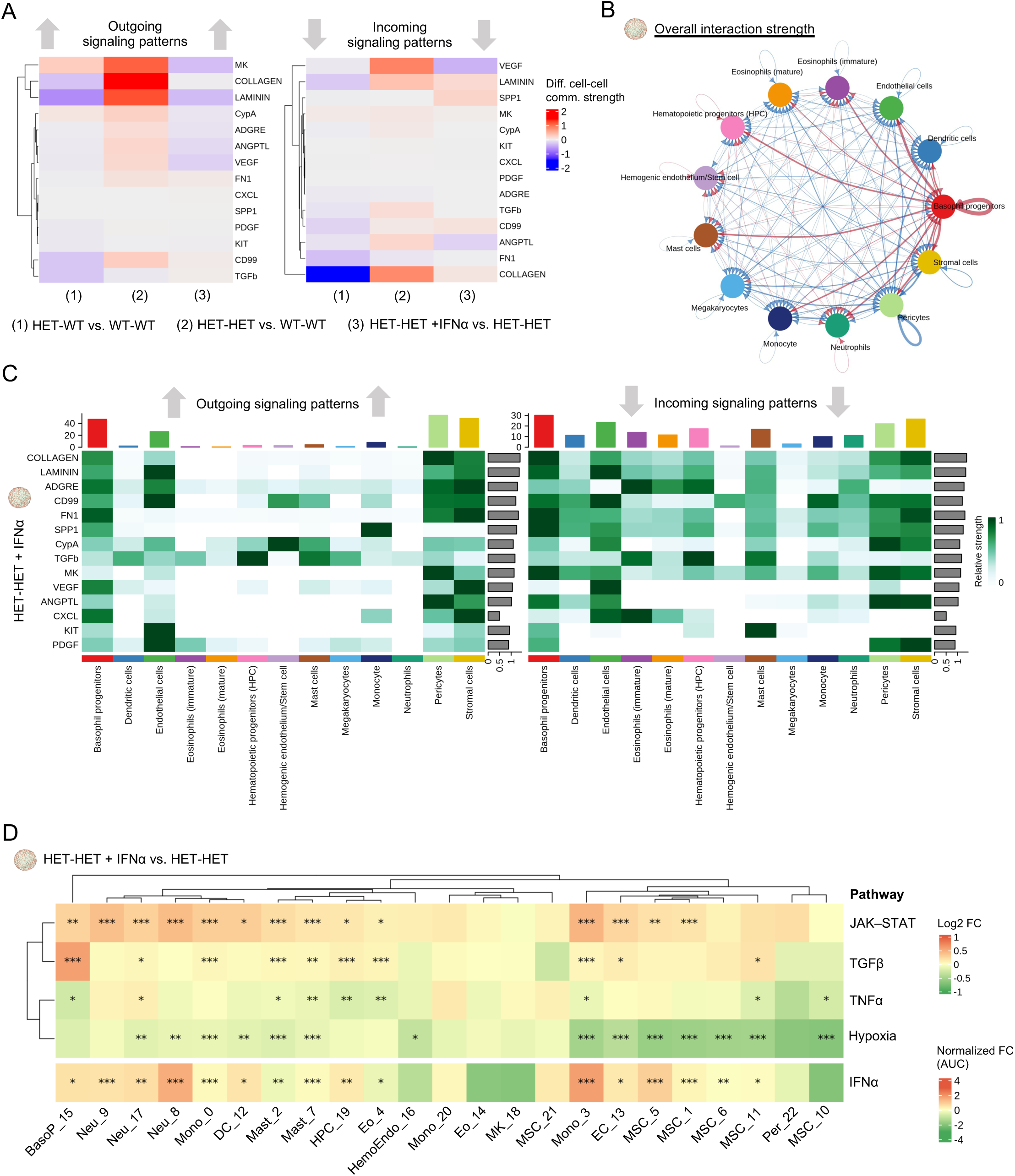
IFNα mitigates pro-fibrotic signaling in JAK2V617F^HET^ assembloids (A) Heatmap showing differential incoming and outgoing signaling in iEC based on L-R analysis from scRNA-seq experiment. The columns represent the tested conditions: (1) HET-WT vs. WT-WT, (2) HET-HET vs. WT-WT, and (3) HET-HET + IFNα vs. HET-HET. Each row corresponds to a mediator of interaction. The color gradient from red to blue indicates increasing to decreasing difference in interaction strength mediated by the listed factors. (B) Circle plot illustrating the overall L-R interaction strength among various cell types in IFNα-treated versus untreated JAK2V617F^HET^ assembloids (HET-HET + IFNα vs. HET-HET). Each node represents a distinct cell type, with MSC clusters categorized as stromal cells. Line thickness denotes interaction strength, while arrows indicate interaction direction. Red lines represent increased interaction strength, whereas blue lines indicate decreased interaction strength. (C) Heatmap illustrating the relative strength of incoming and outgoing signaling among various cell types in JAK2V617F^HET^ assembloids treated with IFNα. Rows represent aggregate strength of signaling for the given pathway, while columns indicate distinct cell types. The color code reflects the strength of the interactions. (D) PROGENy analysis and AUC score analysis of selected pathways in JAK2V617F^HET^ assembloids treated with IFNα (HET-HET + IFNα) compared to untreated JAK2V617F^HET^ assembloids (HET-HET). Abbreviations: FC – Fold change of HET-HET + IFNα vs. HET-HET cocultures.

While IFNα reduced interaction strength across the niche (Fig. 5B), basophil progenitors remained a central signaling hub alongside EC, stromal cells, and pericytes, exhibiting increased signaling via ECM components (collagen, FN1) and growth factors (CXCL, VEGF), but decreased TGFβ-mediated output (Fig. 5C). IFNα also reduced megakaryocyte-derived outgoing TGFβ and PDGF signaling. Additionally, IFNα diminished incoming ECM-related signals (collagen, FN1) to stromal cells and pericytes (Fig. 5C). These changes in outgoing and incoming signaling were accompanied by robust activation of JAK–STAT signaling across hematopoietic lineages. In contrast, CD45⁻ populations exhibited limited responsiveness, with pathway activation largely restricted to MSC clusters 1 and 5 and EC. Reactome pathway analysis confirmed a heterogeneous induction of IFNα target genes across hematopoietic and stromal subsets. However, IFNα consistently attenuated the pro-inflammatory transcriptional signature of JAK2V617F+ assembloids, suppressing hypoxia-related pathways in myeloid and stromal compartments (Fig. 5D).

These findings highlight IFNα’s potential to reverse JAK2V617F-driven endothelial dysfunction and underscore its cell type–specific effects, opening research avenues for mitigating IFNα-associated side effects such as myalgias and fatigue.

### IFNα Inhibits EndMT in Murine Models of PV and MF and in MPN Patients

To validate our findings and assess the translational relevance of our assembloids, we evaluated *in vivo* the effects of IFNα on the BM vasculature using a transgenic JAK2V617F-driven PV mouse model. Eight weeks post-transplantation of JAK2V617F-induced BM cells, pegylated IFNα (pIFNα) or vehicle was administered for 4 or 8 weeks, followed by scRNA-seq analysis of CD31⁺CD45⁻Ter119⁻ BM-derived EC (Fig. 6A; Supplemental Fig. 8A). Vehicle-treated mice developed hallmark PV features, including splenomegaly and elevated white blood cells (WBC), red blood cells, hemoglobin, and neutrophil counts (Fig. 6B–C; Supplemental Fig. 8B), but no increase in BM reticulin deposition, consistent with a PV phenotype (Supplemental Fig. 8C). pIFNα treatment reduced WBC and neutrophil counts and increased Lin⁻Sca1⁺ BM stem and progenitor cells (HSPC), reflecting its known effect on HSPC cycling (Fig. 6C; Supplemental Fig. 8B)^51,52^.

**Figure 6.**
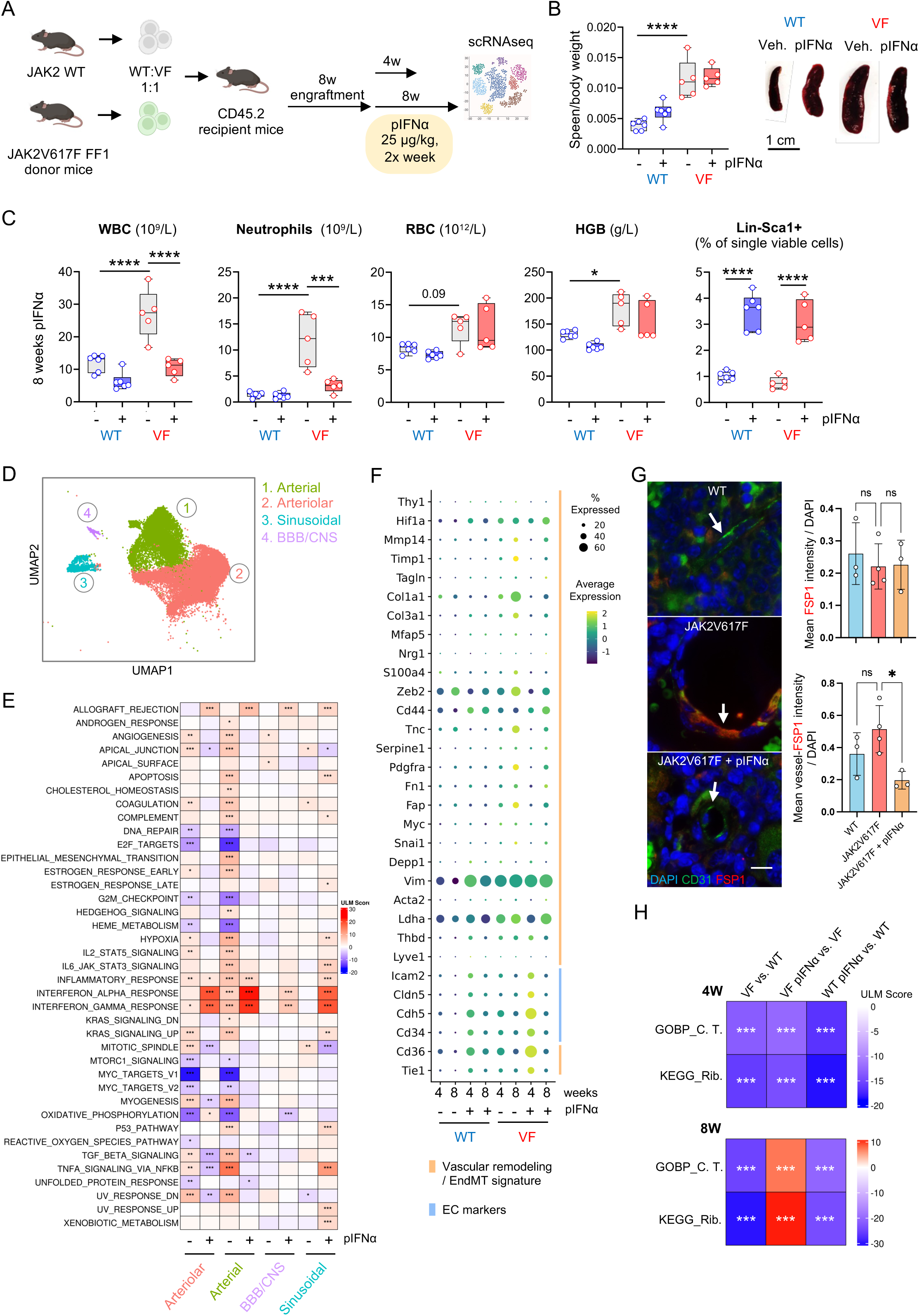
IFNα reverses EndMT and restores endothelial transcriptional programs in a JAK2V617F-driven PV mouse model (A) Schematic overview of the experimental workflow using the transgenic JAK2V617F-driven PV mouse model. Eight weeks post BM transplantation, pIFNα was administered for 4 or 8 weeks. CD31⁺CD45⁻Ter119⁻ BM-derived EC were isolated for scRNA-seq analysis to assess transcriptional changes in response to treatment. (B) Representative images and quantification of spleen size across groups. (C) Peripheral white blood cells (WBC), neutrophil, red blood cells (RBC), and hemoglobin (HGB) levels, and percentage of Lin⁻Sca1⁺ cells in BM from treated and untreated WT and JAK2V617F mice 16 weeks post-engraftment. Two-way ANOVA (n = 5–6). (D) UMAP of CD31⁺CD45⁻Ter119⁻ BM-derived EC clusters identified by scRNAseq. (E) GSEA heatmap showing hallmark pathway enrichment across endothelial subtypes (arteriolar, arterial, BBB/CNS-like, and sinusoidal EC) from scRNA-seq of BM-derived EC 16 weeks post-engraftment. Comparisons: JAK2V617F vs. WT (–), and JAK2V617F + pIFNα vs. JAK2V617F (+). (F) Dot plot showing expression of selected genes across endothelial clusters from WT and JAK2V617F PV mice treated with or without pIFNα for 4 or 8 weeks. Dot size represents the percentage of cells expressing each gene; color scale indicates average expression level (Z-score). Genes are grouped by functional annotation: endothelial markers (blue), and vascular remodeling/EndMT-associated genes (orange). (G) Representative IF of BM sections stained for FSP1 (red) and CD31 (green) in WT and JAK2V617F mice ± pIFNα treatment. Nuclei are counterstained with DAPI (blue). Arrows denote EC. Scale: 10 µm. Quantitative analysis of global and vessel-specific mean FSP1 fluorescence intensity. (H) GSEA for pathways related to cytoplasmic translation (C.T.) and ribosome in endothelial clusters from WT and JAK2V617F mice following 4 weeks (top) and 8 weeks (bottom) of pIFNα treatment. Color scale represents enrichment scores; asterisks indicate statistical significance.

scRNA-seq analysis of CD31⁺CD45⁻Ter119⁻ BM EC identified four endothelial clusters based on canonical marker expression: arterial, arteriolar, sinusoidal, and blood–brain barrier (BBB)/central nervous system (CNS)-like EC (Fig. 6D; Supplemental Fig. 8D–E; Supplemental Table 12). Consistent with their venous identity, sinusoidal EC showed enrichment of the transcription factor *Nr2f2*^53^ and co-expression of the sinusoidal markers *Plvap* and *Stab2*^54,55^.

In diseased mice, GSEA revealed progressive upregulation of pathways linked to vascular remodeling (*angiogenesis*, *apical junction*, *coagulation*, *complement*), inflammation (*hypoxia*, *IL6–JAK–STAT3*, *TNFα–NFκB signaling*, *inflammatory response*), and mesenchymal transition (*epithelial–mesenchymal transition*, *TGF-β signaling*) in arterial EC from weeks 12 to 16, with a similar, albeit attenuated pattern in arteriolar EC. Correspondingly, genes linked to vascular remodeling (*MMP14*, *TIMP1*), ECM production (*Col1a1*, *Col3a1*, *FN1*), EndMT regulation (*Snai1*, *S100a4*), and mesenchymal identity (*FAP*, *PDGFRA*) were progressively upregulated, consistent with an early EndMT phenotype and aligning with our previous *in vitro* findings (Fig. 4B). pIFNα treatment reversed these pathological signatures and downregulated fibrosis-associated ECM genes like *Col1a1*, while inducing robust IFNα/γ responses across all endothelial subsets (Fig. 6E-F; Supplemental Fig. 9A). This transcriptional modulation was supported by immunofluorescence at 12 weeks, showing increased vascular FSP1 deposition in vehicle-treated mice, which was abrogated by pIFNα treatment (Fig. 6G). Consistent with our transcriptomic and TEM findings in JAK2V617F^HOM^ iEC, EC from diseased mice also showed downregulation of ribosome- and translation-related genes (Fig. 6H; Supplemental Fig. 9B). Concomitant suppression of ribosomal genes, E2F targets, and G2/M checkpoint regulators suggests activation of the integrated stress response (ISR)^56^. ISR activation has been implicated in hypoxia tolerance and endothelial dysfunction, both relevant to fibrotic and cardiovascular diseases^57^, but not yet in MPN. These transcriptional dysregulations were reversed after eight weeks of pIFNα treatment (Fig. 6H); however, sinusoidal EC exhibited a comparatively attenuated transcriptional response relative to the arterial and other endothelial subclusters (Supplemental Fig. 9B).

To assess pIFNα’s anti-EndMT effects in the context of BM fibrosis, we employed a TPO-overexpression murine model recapitulating key features of MF. Four weeks after transplantation of TPO-overexpressing KIT⁺ hematopoietic progenitors, mice received pIFNα or vehicle for an additional four weeks (Fig. 7A). Vehicle-treated mice developed characteristic MF features, including splenomegaly, elevated WBC, and increased BM fibrosis (Fig. 7B–C; Supplemental Fig. 10A). pIFNα treatment reduced spleen size and platelet counts and as shown above, increased Lin⁻Sca1⁺ BM HSPC, particularly in empty vector controls (Fig. 7B–D).

**Figure 7.**
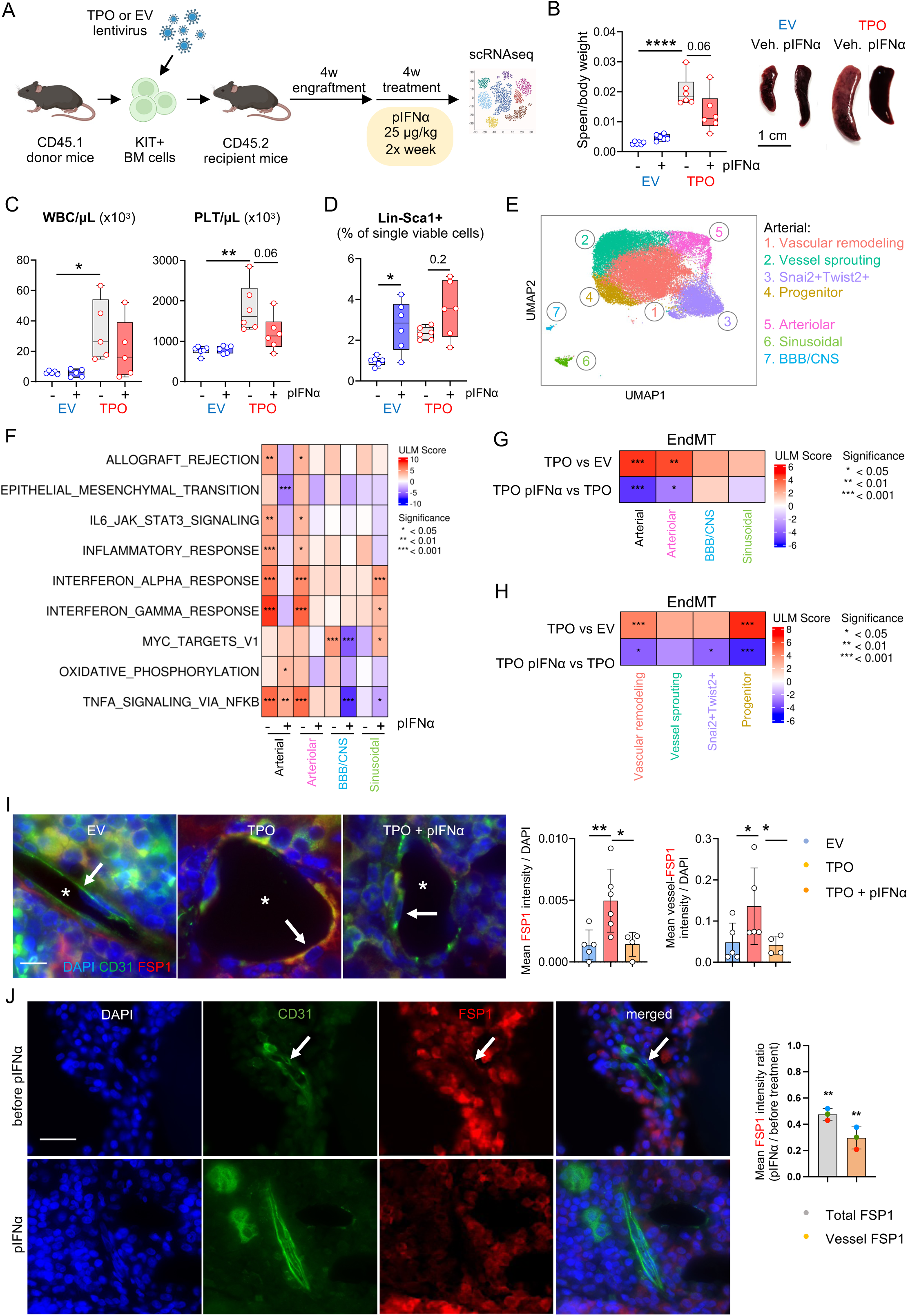
IFNα suppresses EndMT signatures in a murine model of MF (A) Schematic overview of the experimental workflow in the TPO-overexpression MF mouse model. Four weeks after KIT⁺ BM cell transplantation, pIFNα was administered for 4 weeks. CD31⁺CD45⁻Ter119⁻ BM EC were isolated and subjected to scRNA-seq to assess treatment-associated transcriptional changes. (B) Representative spleen images and quantification of spleen size across groups. (C) Peripheral white blood cell (WBC) and platelet (PLT) counts, (D) and percentage of Lin⁻Sca1⁺ progenitor cells in BM. Two-way ANOVA (n = 5). (E) UMAP of CD31⁺CD45⁻Ter119⁻ BM-derived EC clusters identified by scRNAseq. (F) GSEA heatmap displaying pathway enrichment in endothelial clusters. Comparisons: TPO-OE vs. EV (–), and TPO-OE + pIFNα vs. TPO-OE (+). (G) GSEA heatmaps displaying enrichment scores for the curated EndMT signature across endothelial clusters and (H) arterial EC subclusters. (I) Representative IF images of BM sections from EV and TPO-OE mice, treated or not with pIFNα for 4 weeks. Sections were stained for CD31 (green), FSP1 (red), and DAPI (blue). Asterisks denote vessel lumens and arrows indicate EC. Scale: 10 µm. Quantification of global and vessel-specific mean FSP1 fluorescence intensity. Two-way ANOVA (n = 4-6). (J) IF images of BM sections from MPN patients treated with pIFNα, stained for CD31 (green), FSP1 (red), and DAPI (blue). Arrows indicate EC. Scale: 25 µm. Quantification of global and vessel-specific mean FSP1 fluorescence intensity, normalized to baseline. One sample t-test (n = 3). Color indicates individual patients; detailed patient information is provided in the Supplemental Information.

scRNA-seq analysis of CD31⁺CD45⁻Ter119⁻ EC identified the same four EC clusters as in the PV model, with the arterial compartment further subdividing into four transcriptionally distinct subclusters: (1) vascular remodeling, (2) vessel sprouting, (3) Snai2+Twist2+ (inflammation/stress signature) and (4) low-cycling progenitor (Fig. 7E, Supplemental Fig. 10B–C). Consistent with the transcriptional changes observed in the PV model from weeks 12 to 16, GSEA revealed inflammatory activation in arterial and arteriolar EC, including upregulation of TNFα, IL-6–JAK–STAT3, and interferon signaling pathways (Fig. 7F). Interestingly, in both the MF murine model and during disease progression in the PV model, arterial and arteriolar EC showed enrichment of IFNα/γ response–associated genes under basal conditions. While this signature was further amplified by pIFNα treatment in the PV model (Fig. 6E), it was suppressed in the MF model (Fig. 7F). EndMT-associated signatures, including upregulation of ECM-related genes, were observed in arterial (especially subclusters 1 and 4) and arteriolar EC, and were downregulated upon pIFNα treatment (Fig. 7G–H, Supplemental Fig. 10D). Immunofluorescence confirmed reduced FSP1 deposition in BM vessels following pIFNα therapy (Fig. 7I). Analysis of BM biopsies from clinically pIFNα-treated MPN patients confirmed IFNα’s anti-EndMT effect *in vivo* (Fig. 7J).

Together, our findings identify neoplastic arterial remodeling and EndMT as *in vivo* hallmarks of MPN, both reversible by IFNα treatment.

## Discussion

Using patient-specific iPSC, we replicated JAK2V617F-driven neoplastic angiogenesis *in vitro*, a hallmark of BM and spleen remodeling in MPN^2,3^. Drug screening with compounds currently used or under investigation for MPN, revealed toxicity profiles consistent with clinical observations, including vascular side effects of anagrelide^58^ and venetoclax^59^, while ruxolitinib, AMG-232, hydroxyurea, and IFNα demonstrated favorable profiles. pIFNα treatment, approved for PV treatment, has been suggested to improve MF-free survival and preserve near-normal life expectancy in PV^12,60^. However, its mechanisms of action within the malignant clone microenvironment have remained unclear.

RNA-seq revealed zygosity-dependent transcriptional programs in iEC at baseline and following IFNα treatment. JAK2V617F^HOM^ iEC showed suppression of translational pathways and disrupted ribosome-ER organization, while JAK2V617F^HET^ iEC exhibited a pro-mesenchymal phenotype consistent with EndMT, a process implicated in the pathogenesis of MF^4^. EndMT has also been linked to endothelial dysfunction in MPN-associated CVD^6^, a finding particularly relevant in the context of clonal hematopoiesis of indeterminate potential, where JAK2V617F is among the most common mutations and has been directly implicated in vascular pathology^61^. Zygosity-specific alterations in iEC are consistent with previous findings in erythroid cells, where JAK2V617F^HOM^, compared to JAK2V617F^HET^, was associated with enhanced AKT and ERK1/2 signaling^62^. In JAK2V617F^HET^ iEC, dysregulated AKT and ERK1/2 signaling likely promotes EndMT, whereas in JAK2V617F^HOM^ iEC, AKT-driven ROS accumulation^63^ may impair ribosome biogenesis^64^ and trigger heightened ISR, resulting in translational repression^56^. Although hyperproliferation may contribute to ribosomal disorganization in JAK2V617F^HOM^ iEC, IFNα treatment restored translational programs without inducing apoptosis, consistent with its favorable vascular toxicity profile.

In BM niche–mimicking assembloids, we investigated the effects of vascular-restricted JAK2V617F^HET^ expression. Our iPSC-based assembloids supported iEC-MSC pro-angiogenic interactions and recapitulated *in vivo* findings, with acute IFNα exposure enhancing vascularity^65^, and chronic exposure suppressing angiogenesis^14,66^. JAK2V617F^HET^ iEC exhibited EndMT, which was reversed by ruxolitinib, nintedanib, and, to a lesser extent, IFNα. scRNA-seq of genotype-controlled assembloids revealed transcriptional homology to native BM, including stromal heterogeneity and hematopoietic–stromal crosstalk^44^. Vascular JAK2V617F^HET^ expression reduced interaction strength between EC and other niche components, consistent with a vascular environment permissive to JAK2V617F clonal expansion^5^.

Although the mutational burden of the vascular niche *in vivo* remains unclear, recent spatial transcriptomic profiling of BM samples has highlighted vascular niche heterogeneity and revealed activation of pro-inflammatory pathways in MPN patients compared to healthy donors^67^. Our JAK2V617F+ assembloids recapitulated hallmark MPN features, including upregulation of hypoxia-associated genes and TGFβ signaling. The stromal compartment, including pericytes and EC, emerged as a major signaling hub. Consistent with recent findings^50^, basophil progenitors also appeared as central mediators of niche-derived signaling, highlighting their potential as therapeutic targets in MPN. Furthermore, the elevated basal IFNα signaling observed in myeloid subsets is consistent with reports of JAK1/STAT1-driven IFNα sensitization in JAK2V617F+ patients^68^.

Aligning with our *in vitro* findings, in both JAK2V617F-driven PV and TPO-induced MF murine models, EC exhibited pronounced inflammatory signatures (e.g., TNFα signaling via NF-κB), along robust EndMT. This signature intensified between 12- and 16-weeks post-transplantation in the PV model, indicating progressive endothelial dysfunction with disease duration. Concurrently, EC displayed suppression of translation-associated genes, mirroring the transcriptional profile of JAK2V617F^HOM^ iEC *in vitro*.

In the MF model and at advanced disease stages in the PV model, EC showed elevated basal IFNα/γ signaling, suggesting a tonic autocrine IFNα loop in EC^14^, potentially amplified by inflammatory cues during disease progression. This sustained activation may trigger negative feedback mechanisms that reduce treatment response^69^. Consistently, IFNα-responsive genes were upregulated in the PV model following pIFNα treatment, but downregulated in MF, aligning with the reduced clinical efficacy of IFNα in advanced vs. early-stage MPN^70^. Interestingly, this effect was restricted to arterial and arteriolar EC. Sinusoidal EC displayed increased baseline IFNα/γ signaling only in MF and showed a strong transcriptional response to pIFNα, underscoring subtype-specific dynamics within the BM vascular niche. Such heterogeneity may hold therapeutic relevance, particularly in preventing thrombotic complications and JAK2V617F-driven CVD in MPN. The more pronounced transcriptional dysregulation in arterial and arteriolar EC may partly reflect the scRNA-seq enrichment strategy, which favored CD31⁺CD45⁻Ter119⁻ populations and may underrepresent CD31^low^ BM sinusoidal EC^71^.

Across models, IFNα consistently attenuated inflammatory signaling and EndMT-associated transcriptional programs, along with reduced expression of ECM-related genes. The stronger anti-EndMT and anti-fibrotic effects *in vivo* likely reflect prolonged treatment compared to shorter *in vitro* exposure. This aligns with clinical observations identifying IFNα as a rather slow-acting therapeutic^72^. IFNα anti-EndMT potential was also confirmed in patient BM samples, supporting the translational relevance of our findings.

Our findings support a model in which endothelial dysfunction and therapeutic responsiveness of the vascular niche in MPN are shaped by both EC-intrinsic programs and extrinsic inflammatory or clonal cues. Notably, EndMT in JAK2V617F^HET^ iEC was amplified by mutant iHC, while IFNα suppressed EndMT even in assembloids lacking iHC, underscoring endothelial-intrinsic response. These results identify IFNα as a niche-modulating agent capable of cell type-specific transcriptional reprogramming, independent of hematopoietic influence.

While our assembloids offer a valuable platform for dissecting vascular niche dynamics, they do not fully recapitulate BM complexity. Future refinements will improve their translational relevance. Using this model alongside established MPN murine systems, we identify neoplastic arterial remodeling and EndMT as hallmarks of MPN. We further demonstrate that IFNα inhibits EndMT, revealing an unrecognized antifibrotic mechanism that may contribute to its long-term disease-modifying effects and hold therapeutic potential in JAK2V617F-driven CVD. Recent evidence of *in utero* JAK2V617F acquisition^73,74^, suggests that the mutation might occur at the hemangioblast level. In this context, our findings raise the possibility that mutant EC contribute to early niche dysregulation, promoting malignant clonal expansion. Further single-cell and spatial transcriptomic studies are needed to elucidate the vascular mutational burden in MPN patients and to advance the clinical translation of our findings.

## Acknowledgements

This study was supported by the Confocal Microscopy Facility (CMF), the Flow Cytometry Facility (FCF), the Genomics Facility (GF), the Two Photon Imaging Facility (2PIF), and the Immunohistochemistry Facility (IHF), all Core Facilities of the Interdisciplinary Center for Clinical Research (IZKF) Aachen within the Faculty of Medicine at RWTH Aachen University. Transmission Electron Microscopy was performed in the Institute of Pathology and Electron Microscopy Facility, RWTH Aachen. Finally, we would like to thank all the patients who kindly donated samples.

This study was in part funded by grants to Marcelo A. S. de Toledo from the RWTH Aachen University START grant (128/22), to Steffen Koschmieder from the German Research Foundation (Deutsche Forschungsgemeinschaft, DFG) (KO 2155/6-1, project number AOBJ 636363; and KO2155/8-2, project number AOBJ: 680695) and by funds from German Research Foundation as part of the Clinical Research Unit CRU 344 to SK, NC, IC, and THB (KO2155/7-1, CH1509/1-1, GE2811/4-1 and BR1782/5-1). Madeline J. Caduc was funded by the Clinician Scientist Program Junior, the “FF-Med” funding program for female doctors in the post-doc phase of the Faculty of Medicine, RWTH Aachen, Germany and the Mildred-Scheel-Postdoc Program of the German Cancer Aid (Deutsche Krebshilfe, DKH).

## Author Contributions

**M. J. C.:** conceptualization, formal analysis, investigation, visualization, methodology, manuscript–original draft, writing–review and editing. **M. H. E. M.:** methodology. **M. G.:** methodology. **P. W.:** methodology. **K.G.:** methodology. **J. E. P.:** methodology**. D. W. L. W.:** methodology, visualization. **C. B. L.:** methodology. **L. B.:** methodology. **P. S.:** methodology. **B. J.:** methodology. **E. M. B.:** methodology, visualization. **S. E.:** methodology, visualization. **G. M-N.:** resources, investigation, writing-review, and editing. **M. V.:** methodology, visualization. **J. E.:** resources. **P. B.:** resources, investigation, methodology. **M. A.:** resources, investigation, methodology. **N. C.:** resources, writing-review, and editing. **J. F.:** methodology, writing-review, and editing. **I. G. C.:** resources, methodology. **M. Z.:** resources, investigation, writing-review, and editing. **R. K. S.:** resources, writing-review, and editing**. R. S.:** resources, writing-review, and editing**. T. H. B.:** resources, funding acquisition, writing-review, and editing. **S. K.:** conceptualization, supervision, resources, funding acquisition, project administration, investigation, data analysis, writing-review, and editing. **M. A. S. T.:** conceptualization, supervision, resources, funding acquisition, project administration, investigation, methodology, formal analysis, writing-review, and editing.

---

**Supplemental Fig. 1.**
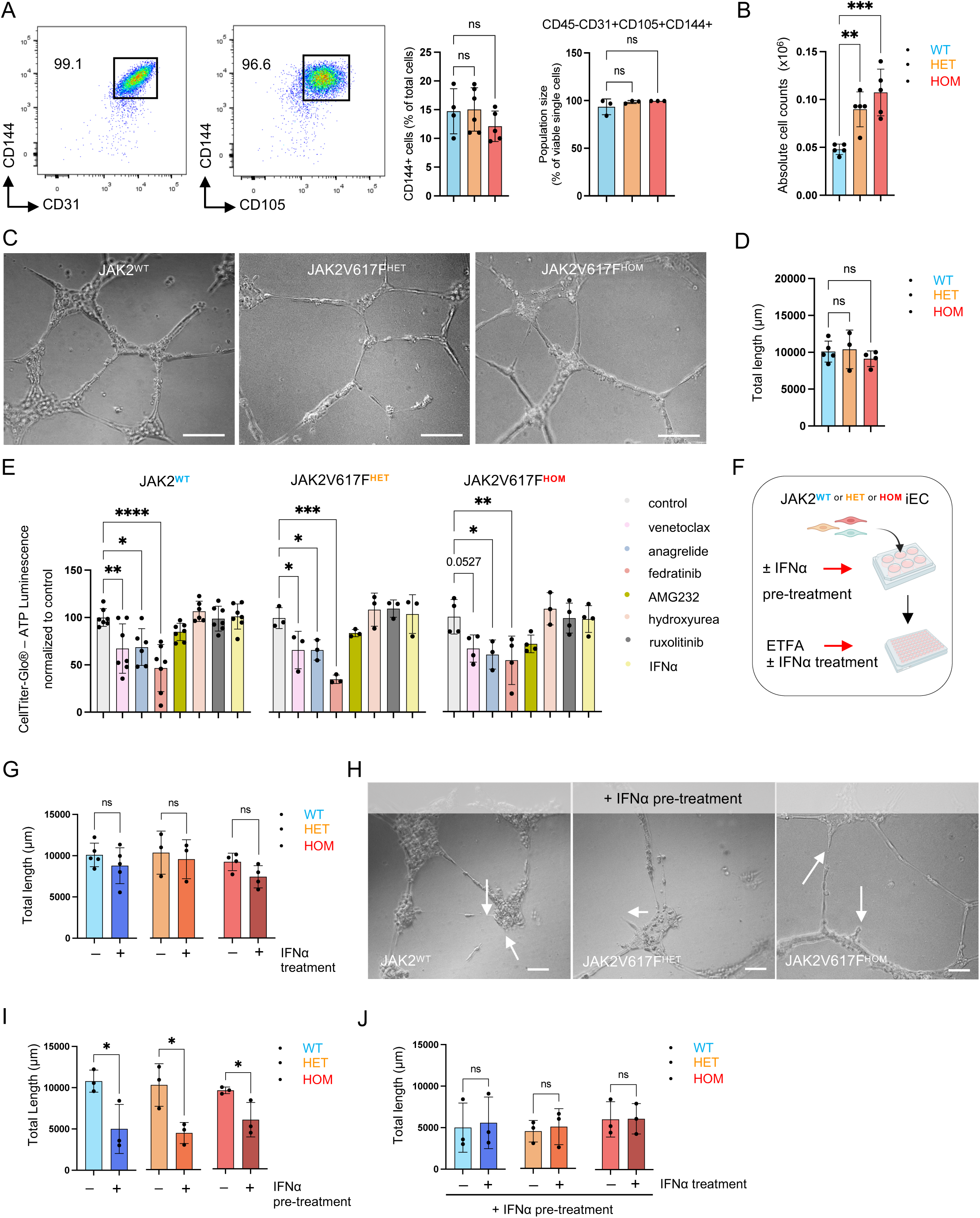

**Supplemental Fig. 2.**
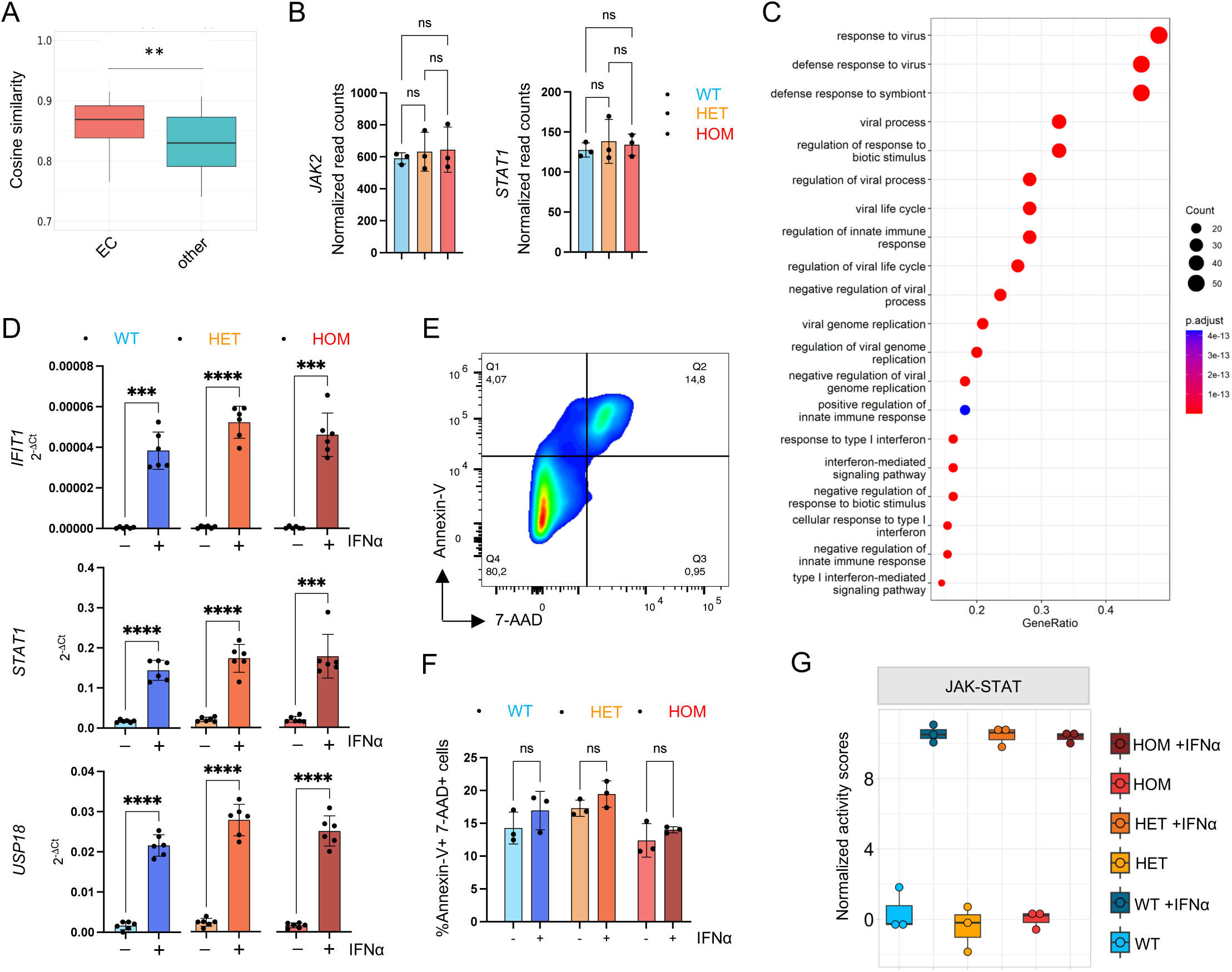

**Supplemental Fig. 3.**
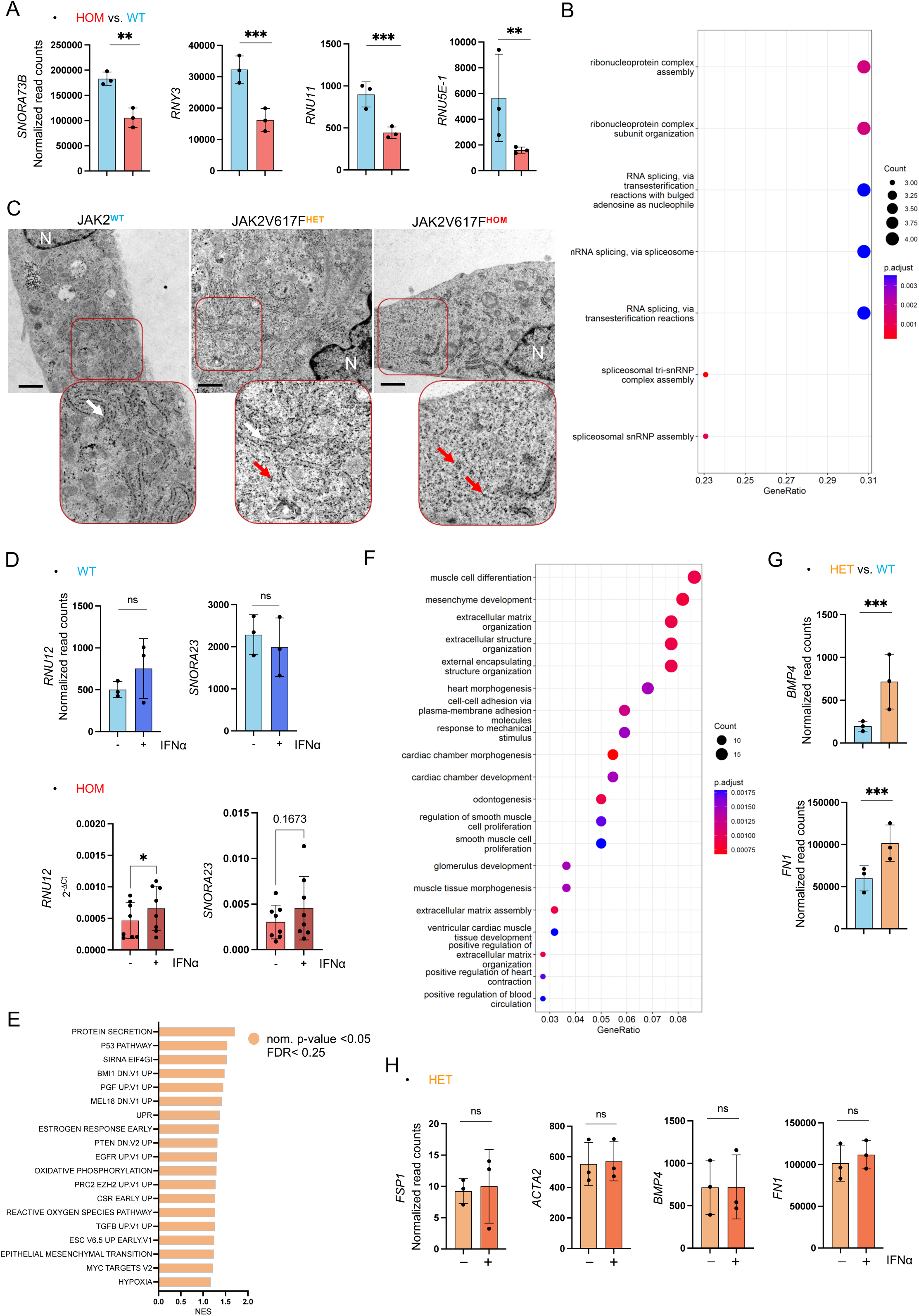

**Supplemental Fig. 4.**
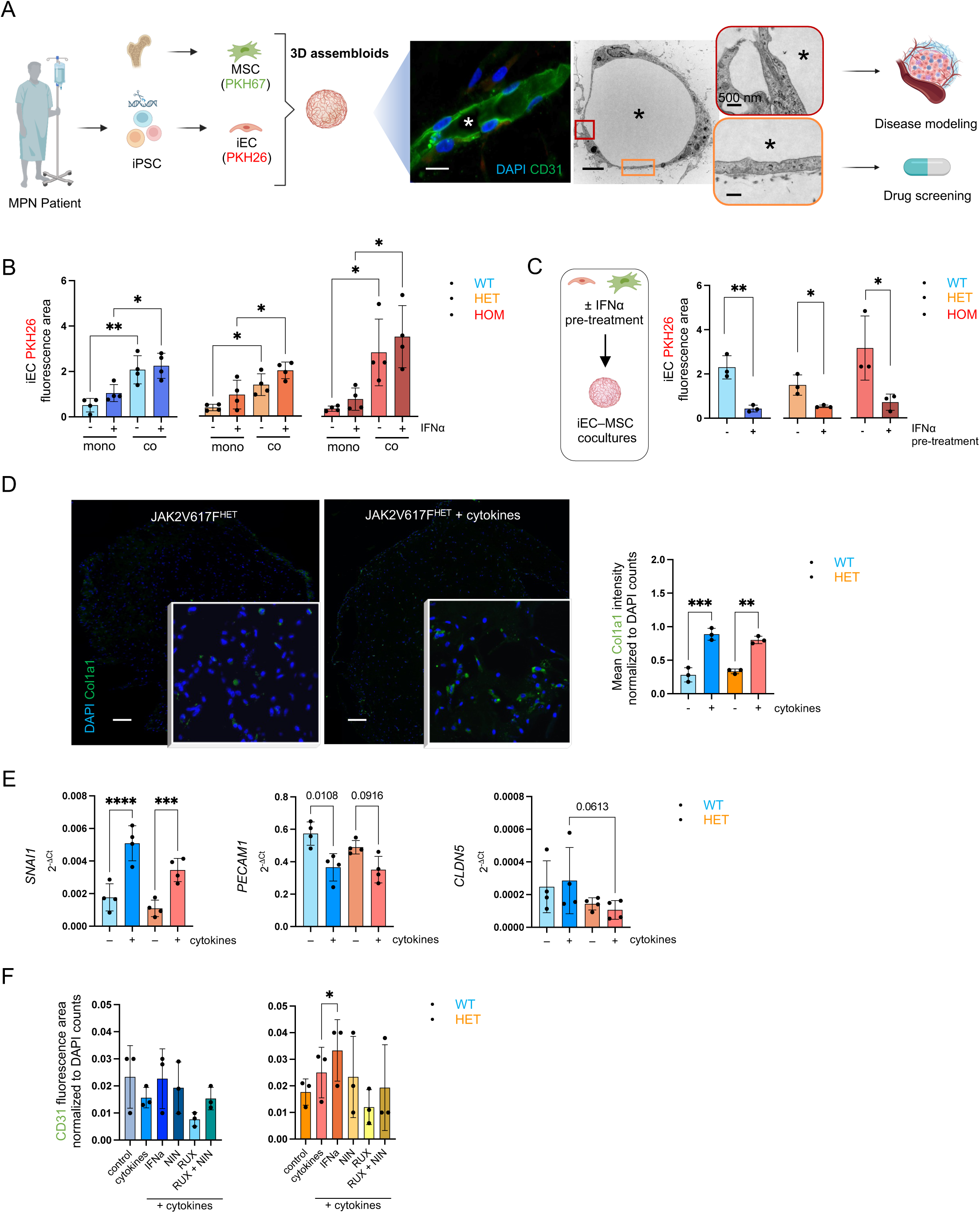

**Supplemental Fig. 5.**
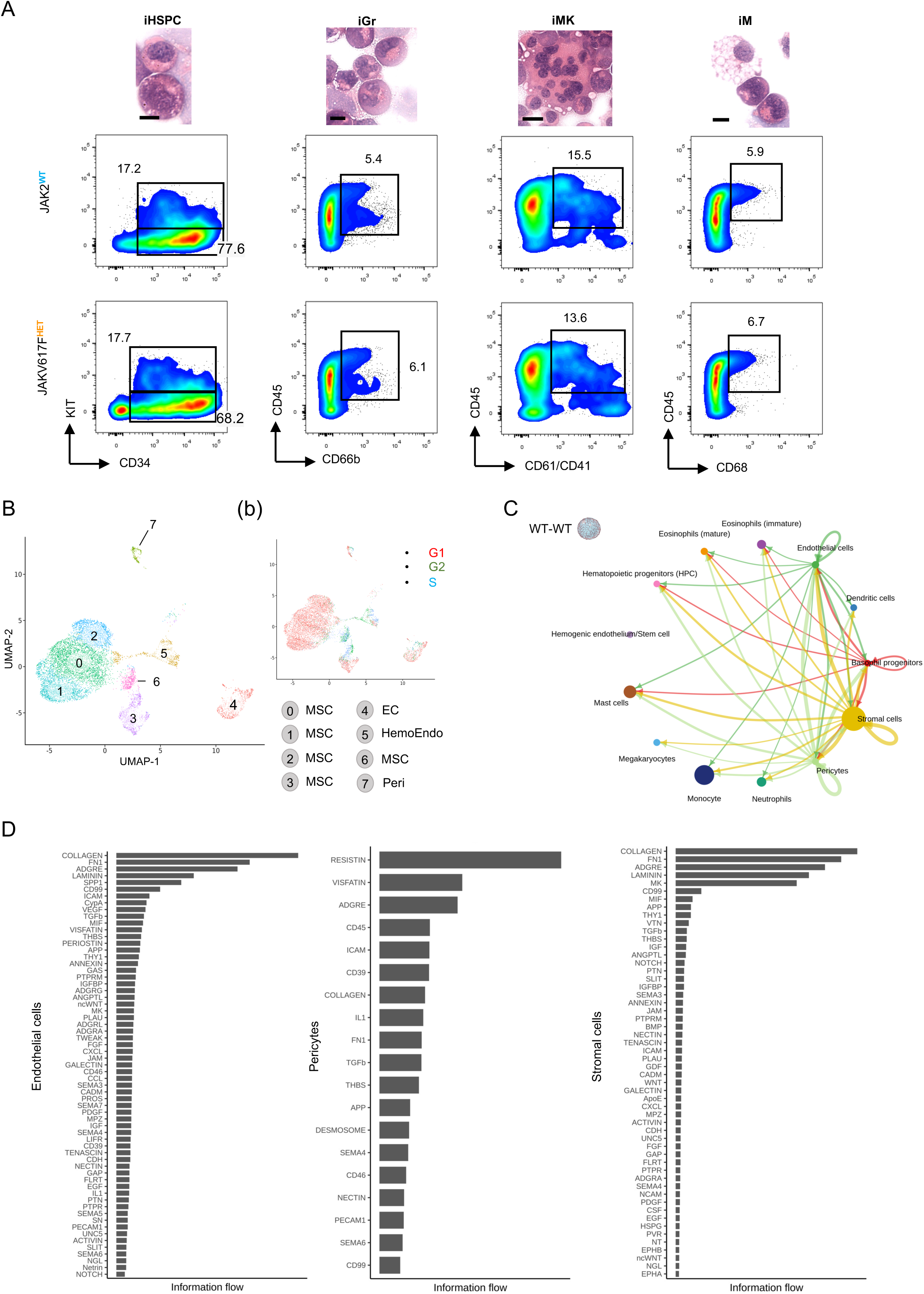

**Supplemental Fig. 6.**
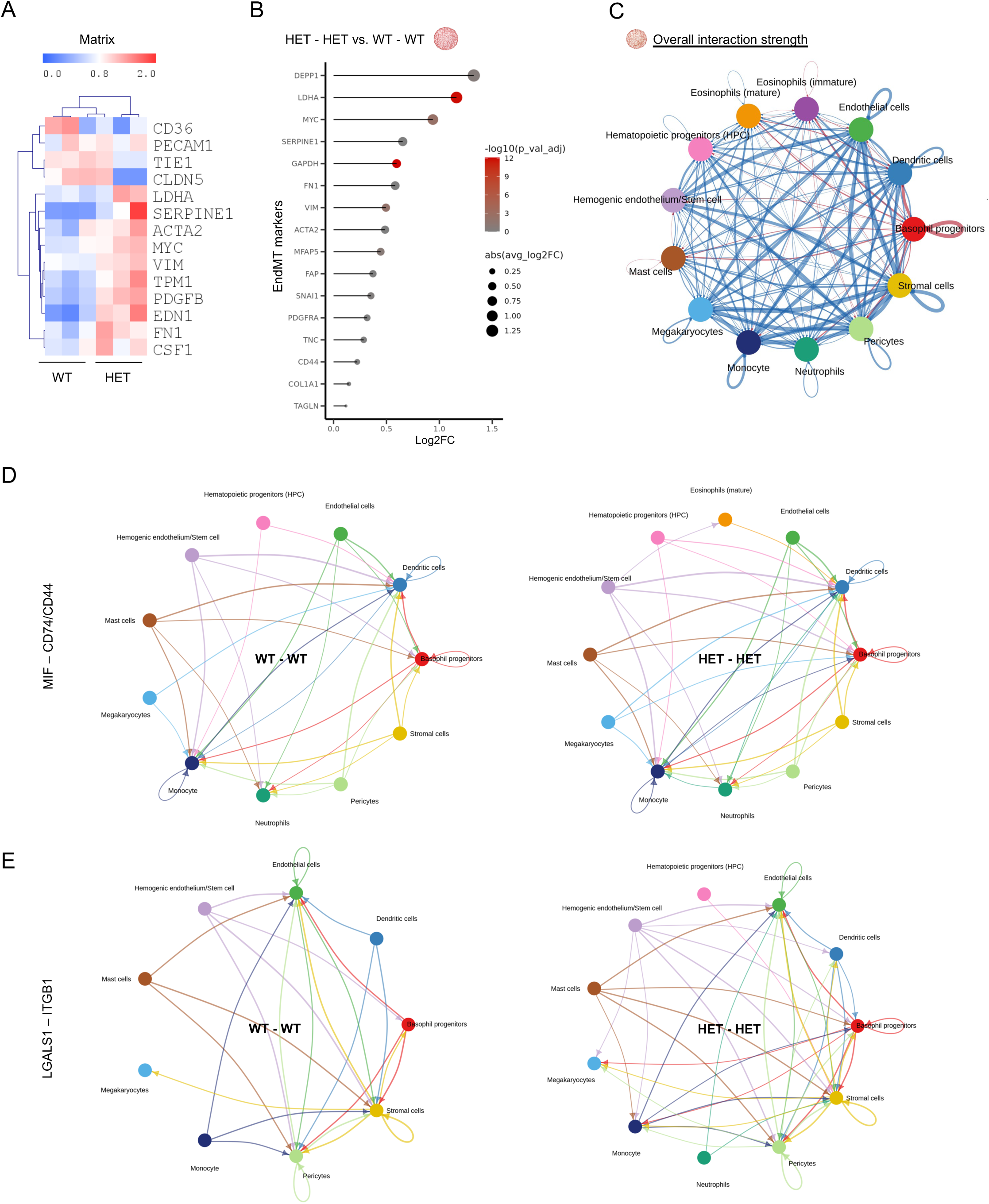

**Supplemental Fig. 7.**
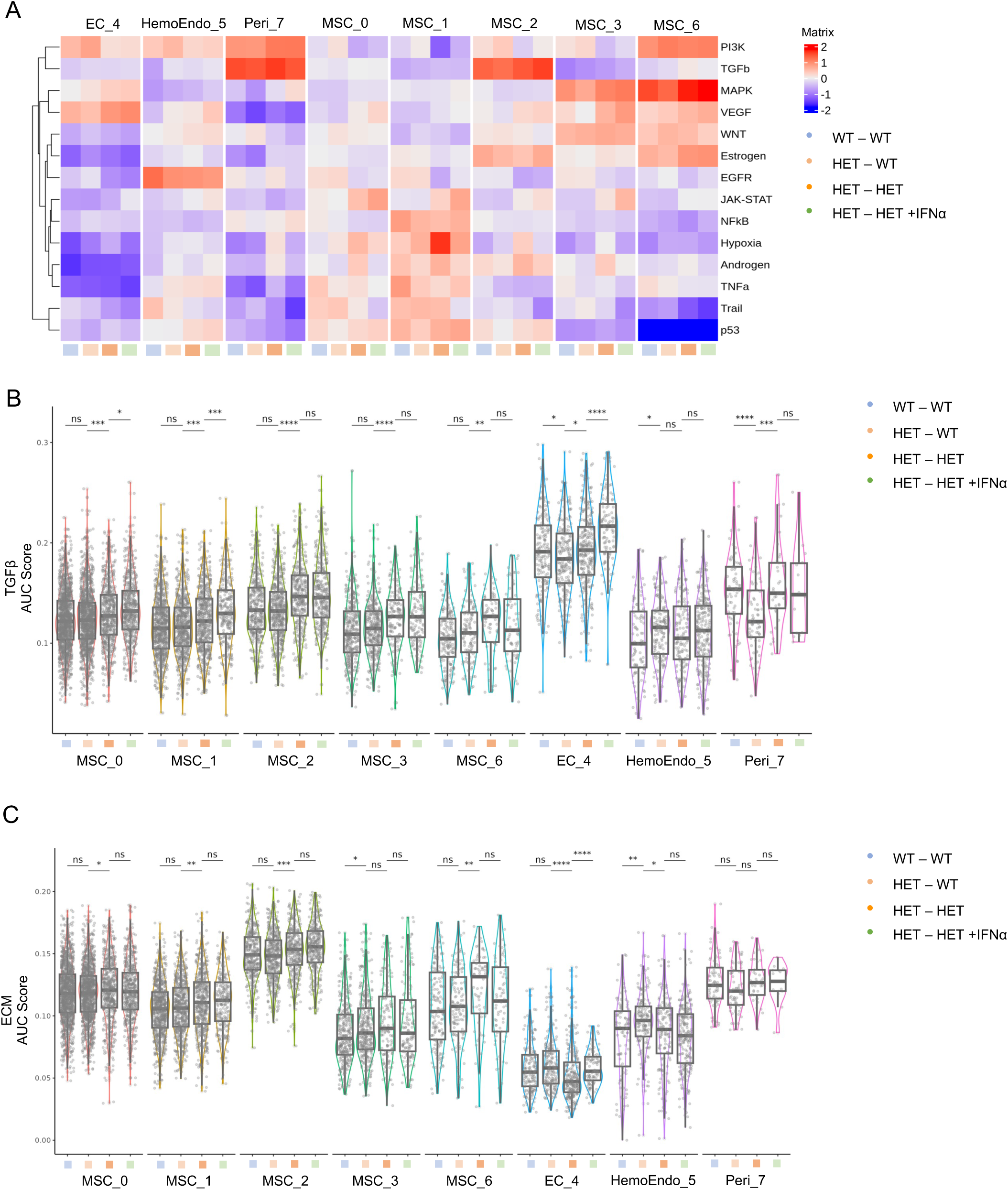

**Supplemental Fig. 8.**
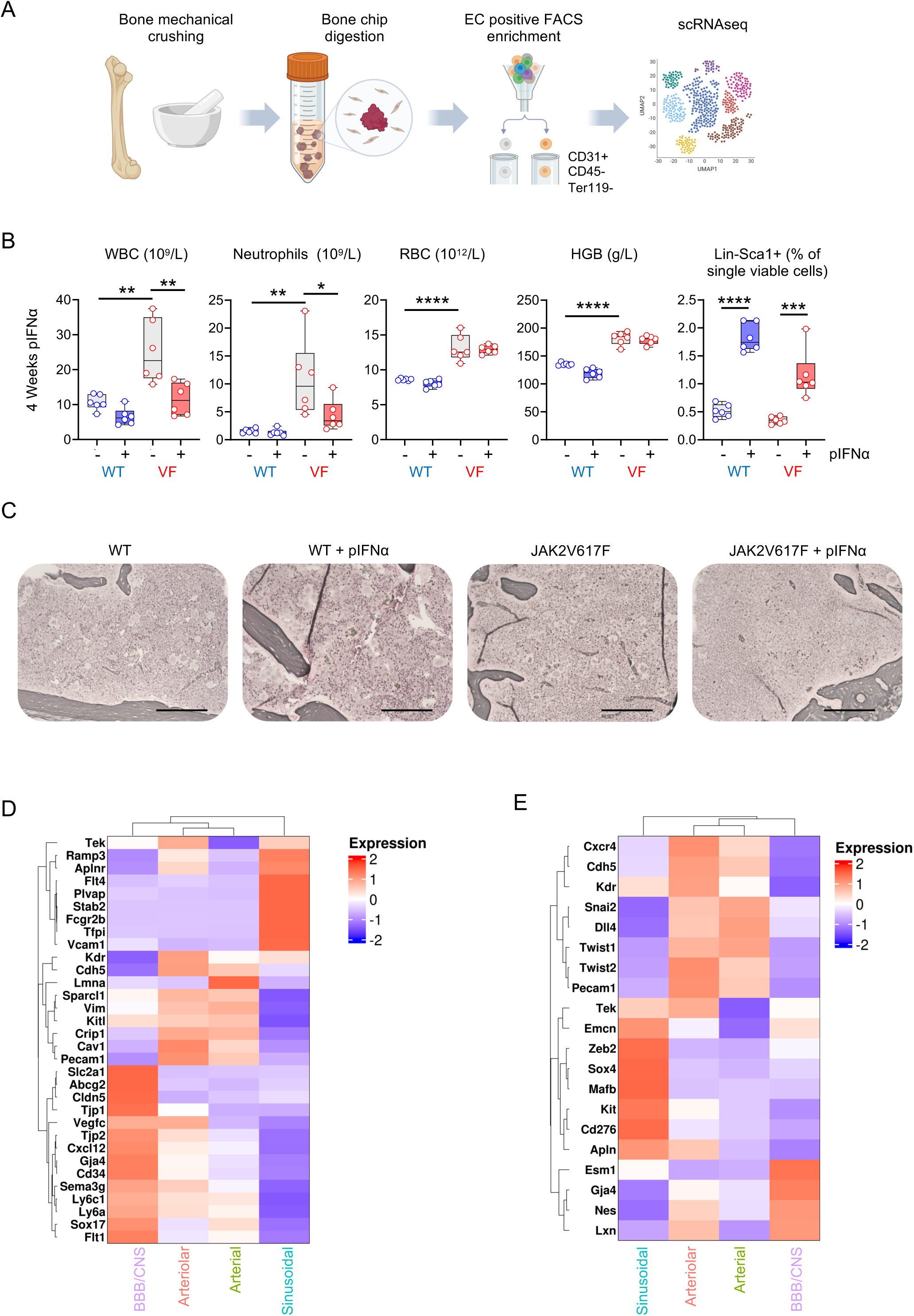

**Supplemental Fig. 9.**
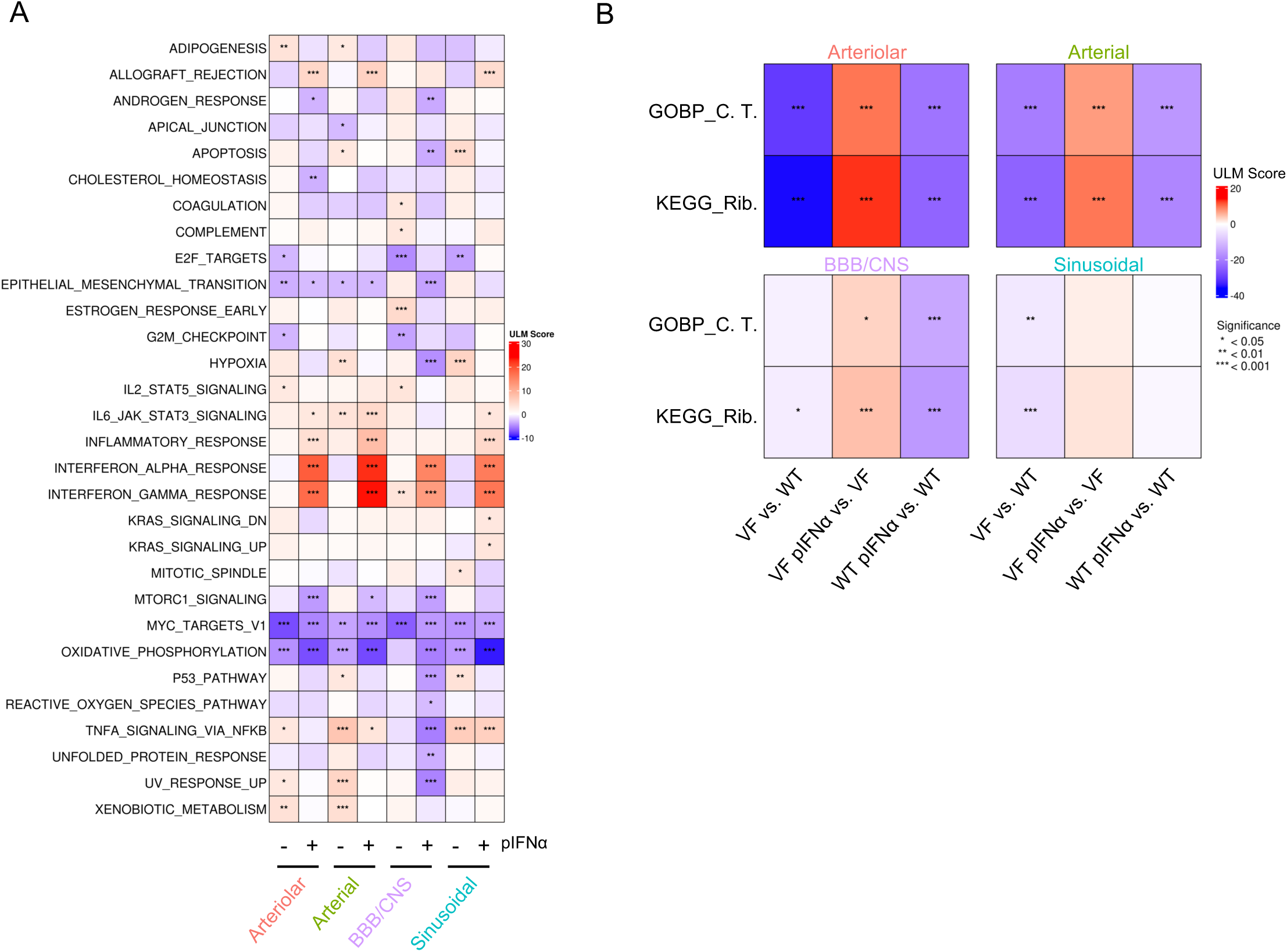

**Supplemental Fig. 10.**
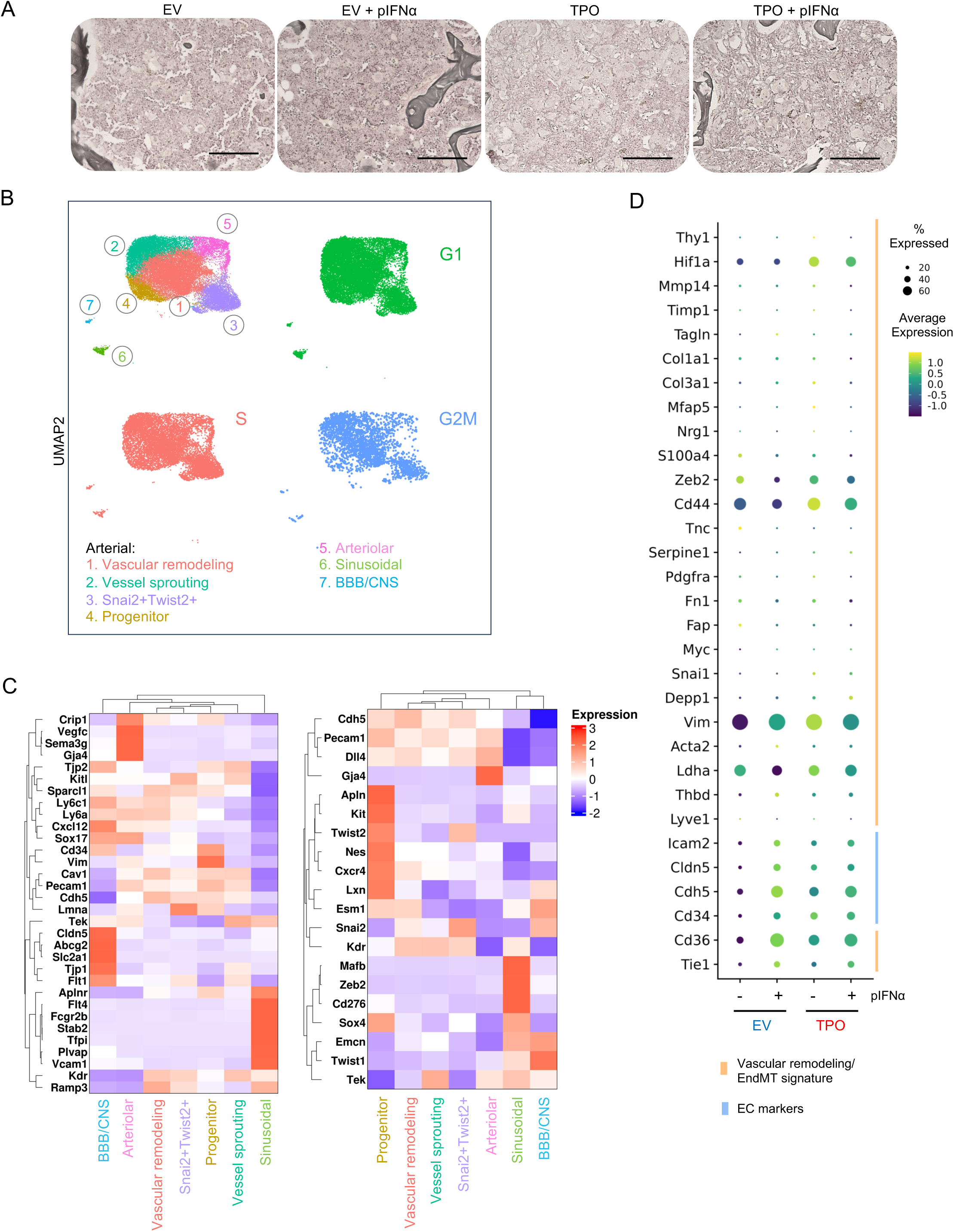

