## Supplemental Methods for "Inhibition of JAK2V617F-Driven Neoplastic Endothelial Dysfunction by IFNα as a Novel Antifibrotic Mechanism"

**Supplemental Table 1**. List of iPSC clones

| Patient | Clone name | Genotype | Generated by |
|---|---|---|---|
| Patient 1 | PV1 007 WT | $JAK2^{WT}$ | Reprogramming |
| Patient 1 | PV1 009 HET | $JAK2V617F^{HET}$ | Reprogramming |
| Patient 1 | PV1 012 HOM | $JAK2V617F^{HOM}$ | CRISPR/Cas9 |
| Patient 1 | PV1 033 HOM | $JAK2V617F^{HOM}$ | CRISPR/Cas9 |
| Patient 2 | PV2 072 WT | $JAK2^{WT}$ | CRISPR/Cas9 |
| Patient 2 | PV2 092 WT | $JAK2^{WT}$ | CRISPR/Cas9 |
| Patient 2 | PV2 070 HET | $JAK2V617F^{HET}$ | CRISPR/Cas9 |
| Patient 2 | PV2 021 HOM | $JAK2V617F^{HOM}$ | Reprogramming |

**Supplemental Table 2**. Fibrin hydrogels composition

| | Final concentration | Ratio | Company |
|---|---|---|---|
| Fibrinogen | 5 mg/ml | 50% | Merck |
| Thrombin | 2,5 U/ml | 20% | Sigma Aldrich |
| EBM-2 | | 25% | Lonza |
| Aprotinin | 500 µg/ml | 5% | Sigma Aldrich |

**Supplemental Table 3**. List of antibodies used for IF

| Primary antibody | Dilution | Reference |
|---|---|---|
| CD31 anti-human | 1:100 | Invitrogen |
| CD31 anti-mouse | 1:100 | Dianova |
| CD31 anti-human | 1:100 | MA5-33063, ThermoFisher |
| FSP1 anti-human, mouse | 1:100 | 07-2274, Millipore |
| FSP1 anti-human, mouse | 1:100 | Ab93283, Abcam |
| Collagen1a1 anti-human | 1:150 | MAB6220, R&D Systems |
| αSMA anti-human, mouse, rat | 1:500 | MAB1420, R&D Systems |
| **Secondary antibody** | | |
| Alexa Fluor 750 goat anti-rabbit IgG (H+L) | 1:200 | A21039, ThermoFisher |
| Alexa Fluor 488 goat anti-mouse IgG (H+L) | 1:200 | A28175, ThermoFisher |
| Alexa Fluor® 647 AniPure Goat Anti-Rat IgG (H+L) | 1:100 | AB_2338393, Jackson ImmunoResearch Europe |

**Supplemental Table 4.** Oligonucleotides used for RT-PCR analysis (5'–3')

| | |
|---|---|
| SNAI1 | FRW CATGTCCGGACCCACACTG<br>REV GACATTCGGGAGAAGGTCCG |
| FSP1 | FRW GGAGACAGGGTTCGCCAAAA<br>REV CCCGTGGCCAGGATAAGATG |
| ACTA2 | FRW TCCTTCATCGGGATGGAGTCT<br>REV TACATAGTGGTGCCCCCTGA |
| RNU12 | FRW AGACTGACTGTGGGGTGGTC |

| | |
|---|---|
| | REV GTGGGTCCCAACGTCAATAC |
| SNORA23 | FRW TGGTAGCAGTGTCTGTCTGTG<br>REV CCAAGTTACTCTTTGGCCGC |
| PECAM1 | FRW GAGTCCTGCTGACCCTTCTG<br>REV ATTTTGCACCGTCCAGTCC |
| CLDN5 | FRW ACCTGACAGATACACGCTCG<br>REV CCCTCTTTGAAGGTTCGGGG |
| IFIT1 | FRW CTCCCTAATTTACAGCAACCATGA<br>REV ACCATTTGTACACATCTCCACTGT |
| STAT1 | FRW TGTATGCCATCCTCGAGAGC<br>REV AGACATCCTGCCACCTTGTG |
| USP18 | FRW ATCACGAATGAGCAAATTTGG<br>REV CAACCAGGCCATGAGGGTAG |

**Supplemental Table 5.** List of FACS antibodies

| **Anti-human** | **Company** |
|---|---|
| CD31-PE - clone WM59 | BD Bioscience |
| CD34-APC - clone 581 BD | Bioscience |
| CD45-APC-Cy7 - REA747 | Miltenyi Biotec |
| CD61-FITC - clone VI-PL2 | Biolegend |
| CD66b-PE - clone REA306 | Miltenyi Biotec |
| CD105-FITC – clone 43A4E1 | Miltenyi Biotec |
| CD117 (KIT)-PE-Cy7 - clone104D2 | eBioscience |
| CD144-FITC – clone REA199 | Miltenyi Biotec |
| **Anti-mouse** | |
| c-kit-APC-Cy7 | Biolegend |
| CD150-APC | Biolegend |
| CD48-PB | Biolegend |
| Sca1-PE-Cy7 | Biolegend |
| CD11b-PE-Cy5 | Biolegend |
| Gr-1-PE-Cy5 | Biolegend |
| CD4-PE-Cy5 | Biolegend |
| CD8-PE-Cy5 | Biolegend |
| CD3-PE-Cy5 | Biolegend |
| Ter119-PE-Cy5 | Biolegend |

**Supplemental Table 6**. List of MPN-/ anti-fibrotic therapeutics tested on iEC

| **Name** | **Company** | **Reference** |
|---|---|---|
| Anagrelide | Selleckchem | S3172 |
| Fedratinib | Selleckchem | S2736 |
| Hydroxyurea | Selleckchem | S1896 |
| AMG 232 (Navtemadlin) | Selleckchem | S7426 |

| | | |
|---|---|---|
| Nintedanib | Selleckchem | S1010 |
| Rh IFN-alpha 2b | Immunotools | 11343516 |
| Ruxolitinib | Selleckchem | S1378 |
| Venetoclax | Selleckchem | S8048 |

**Supplemental Table 7**. Overview of iEC samples used for global RNAseq

| Batch | iEC clone | Tested condition |
|---|---|---|
| 1 | WT 007 | Control |
| 1 | WT 007 | + IFNα |
| 1 | WT 072 | Control |
| 1 | WT 072 | + IFNα |
| 1 | WT 092 | Control |
| 1 | WT 092 | + IFNα |
| 1 | HET 009 | Control |
| 1 | HET 009 | + IFNα |
| 1 | HOM 021 | Control |
| 1 | HOM 021 | + IFNα |
| 2 | HET 009 | Control |
| 2 | HET 009 | + IFNα |
| 2 | HOM 021 | Control |
| 2 | HOM 021 | + IFNα |
| 3 | HOM 021 | Control |
| 3 | HOM 021 | + IFNα |
| 4 | HET 009 | Control |
| 4 | HET 009 | + IFNα |

**Supplemental Table 8.** Cosine similarity of iEC compared to reference EC

| Reference sample | iPSC |
|---|---|
| Adrenal-Vascular endothelial cells | 0.9115 |
| Cerebellum-Vascular endothelial cells | 0.907 |
| Cerebrum-Vascular endothelial cells | 0.8886 |
| Eye-Vascular endothelial cells | 0.7872 |
| Heart-Lymphatic endothelial cells | 0.838 |
| Heart-Vascular endothelial cells | 0.8784 |
| Kidney-Vascular endothelial cells | 0.8817 |
| Liver-Vascular endothelial cells | 0.8409 |
| Lung-Lymphatic endothelial cells | 0.88 |
| Lung-Vascular endothelial cells | 0.9008 |
| Muscle-Vascular endothelial cells | 0.8594 |
| Pancreas-Lymphatic endothelial cells | 0.8385 |
| Pancreas-Vascular endothelial cells | 0.9153 |

| | |
|---|---|
| Spleen-Vascular endothelial cells | 0.8395 |
| Stomach-Lymphatic endothelial cells | 0.7657 |
| Stomach-Vascular endothelial cells | 0.8022 |

**Supplemental Table 9.** Qualitative ultrastructural analysis of iEC via TEM

| **iPSC clone** | **ER amount** | **Ribosome amount** | **Structure clearness** | **Golgi amount** |
|---|---|---|---|---|
| HOM 012 | ++ | ++ | + | + |
| HOM 021 | + | ++ | + | ++ |
| HOM 033 | + | + | + | + |
| HET 070 | ++ | ++ | +++ | + |
| HET 009 | ++ | + | + | ++ |
| WT 092 | ++ | + | +++ | ++ |
| WT 007 | + | + | ++ | ++ |

**Supplemental Table 10.** List of genotype-specific DEG following IFNα treatment

**Supplemental Table 11.** Gene list for cell cluster annotation in scRNA-seq of 3D assembloids

**Supplemental Table 12.** Marker genes used for endothelial cluster annotation in scRNA-seq analysis of MPN murine models

### Supplemental Methods

#### Differentiation of iPSC into Endothelial Cells

iPSC were differentiated towards endothelial cells (EC) as previously described[1,2]. Briefly, iPSC were dissociated with accutase (StemCell Technologies) and seeded on Matrigel-coated 6-well plates at a density of $5x10^5$ cells/well in StemMACS iPS-Brew XF (Miltenyi Biotec) containing 10µM Rho-Kinase inhibitor (Abcam). The next day, start of the differentiation protocol with cultivation of the cells in STEMdiff Mesoderm Induction Medium (StemCell Technologies) until day 5. From day 6 to 8, medium was switched to StemPro-34 SFM (Thermo Fisher Scientific) supplemented with 200 ng/ml VEGF (Peprotech) and 2µM Forskolin (Sigma-Aldrich). From day 9 to 12, cells were cultivated in StemPro-34 SFM supplemented with 50ng/mL VEGF. Medium change was performed daily. Cells were then dissociated using Accutase and selected for CD144 surface expression by Magnetic-activated cell sorting (MACS) (Miltenyi Biotec). The CD144+-enriched cells were then maintained on 0,1% gelatin in Endothelial Cell Growth Medium-2 (EGM-2, Lonza). Endothelial differentiation was monitored by flow cytometry (FACS) with antibodies specific for CD31, CD34, CD45, CD105 and CD144 (Supplemental Table 5).

#### Hematopoietic Differentiation of iPSC via Spin-EB Protocol

Human iPSC were differentiated into hematopoietic cells (iHC) as previously described[3,4]. Briefly, 3,000 iPSC per well were seeded into U-bottom 96-well plates in cytokine-supplemented serum-free medium (SFM). Embryoid body (EB) formation was induced by centrifugation at 380 × *g* for 5 minutes. From day 2 to day 8, EB were cultured in SFM supplemented with VEGF (10 ng/mL), BMP4 (10 ng/mL; Miltenyi Biotec), bFGF (10 ng/mL; Peprotech), and SCF (Miltenyi Biotec). BMP4 and VEGF were withdrawn from the medium starting on day 8.

From day 10 onward, lineage-specific differentiation was promoted using the following cytokine combinations:

- Megakaryocytic lineage: SCF (100 ng/mL) + TPO (40 ng/mL)
- Granulocytic lineage: SCF (100 ng/mL) + IL-3 (30 ng/mL) + G-CSF (100 ng/mL)
- Monocytic/macrophage lineage: SCF (100 ng/mL) + IL-3 (30 ng/mL) + M-CSF (100 ng/mL) + FLT3L (50 ng/mL)

On day 14, cells were harvested and passed through a 100 µm cell strainer to separate EB from the non-adherent suspension cells. The resulting single-cell suspension was subsequently analyzed by flow cytometry using antibodies against CD45, CD34, CD41/CD61, CD66b, CD68, and CD117.

**Cell Morphology Analysis**

For morphological characterization of iEC and iHC, single-cell suspensions were cytocentrifuged onto glass slides using a Shandon Cytospin 4 cytocentrifuge (Thermo Fisher Scientific). Cells were fixed in methanol and stained with hematoxylin and eosin (H&E; Sigma-Aldrich). Slides were mounted using Entellan (Merck), and images were acquired with an EVOS M7000 Cell Imaging System (Thermo Fisher Scientific).

**Flow Cytometry Analysis**

Following harvesting, cells were washed with ice-cold FACS buffer (PBS containing 2 mM EDTA and 2% bovine serum albumin) and incubated with lineage-specific antibodies for 30 minutes at 4 °C. A complete list of antibodies used is provided in Supplemental Table 5. Stained and unstained samples were analyzed using a FACS Canto II flow cytometer (BD Biosciences), and data were processed using FlowJo software (BD Biosciences).

**Proliferation Assay**

On day 0, iEC were seeded in triplicate into 24-well plates at a density of 20,000 cells per well and cultured in supplemented EGM-2 medium with daily medium changes. On day 4, cells were harvested using Accutase, and total cell numbers were determined manually using a Neubauer hemocytometer.

**Drug Screening Using MPN Patient-Derived iEC**

iEC were seeded at a density of 20,000 cells per well into 96-well plates pre-coated with 0.1% gelatin and cultured in supplemented EGM-2. On day 1, half-medium change was performed. On day 2, candidate therapeutics (listed in Supplemental Table 6) were added to freshly renewed EGM-2 at a final concentration of 1 µM (IFNα: 1000 U/mL). Cells were incubated in 100 µL of drug-containing medium for 48 hours without further medium exchange. Negative controls were maintained in EGM-2 alone.

Cell viability was assessed using the CellTiter-Glo Luminescent Cell Viability Assay (Promega) on a SpectraMax i3 Plate Reader (Molecular Devices) and analyzed with SoftMax Pro software. Luminescence values were normalized to the corresponding untreated control for each genotype. All treatment conditions were performed in triplicate, and each genotype was tested in at least three independent experiments ($JAK2^{WT}$ n=6, $JAK2V617F^{HET}$ n=3, $JAK2V617F^{HOM}$ n= 3-4).

**Endothelial Tube Formation Assay**

To assess the anti-angiogenic effects of IFNα, endothelial tube formation assays (ETFA) were performed in the presence or absence of IFNα (1000 U/mL), administered either during the starvation phase (pre-treatment), during the incubation on Matrigel/Geltrex (treatment), or both. $JAK2^{WT}$ and JAK2V617F iEC were cultured to confluency and starved for 4 hours in Endothelial Basal Medium-2 (EBM-2) supplemented with 2% FCS prior to harvesting. Cells were then seeded at a density of 20,000 cells per well into 96-well plates pre-coated with either Matrigel (Corning) or Geltrex (Gibco), using EBM-2 medium supplemented with 2% FCS and 25 ng/mL VEGF. After 12–14 hours of incubation, four images per well were acquired using an EVOS Cell Imaging System. Quantification of tubule-like structures was performed using the Angiogenesis Analyzer plugin for ImageJ developed by Carpentier *et al.* [5].

**RNA Extraction and RT-PCR**

Total RNA iEC was extracted using the RNeasy Mini Kit (Qiagen) according to the manufacturer's instructions. RNA concentration and purity were assessed using a NanoDrop spectrophotometer (Thermo Fisher Scientific). Complementary DNA (cDNA) synthesis was performed using M-MLV reverse transcriptase (Invitrogen). Quantitative real-time PCR (RT-qPCR) was conducted on a 7500 Fast Real-Time PCR System (Applied Biosystems, Life Technologies) using SYBR Select Master Mix (Thermo Fisher Scientific). Primers were obtained from Eurofins-MWG Biotech (Ebersberg, Germany) and are listed in Supplemental Table 4. Gene expression levels were normalized to GAPDH, and relative mRNA expression was calculated using the $2^{-\Delta CT}$ method.

**Global RNA-sequencing of iEC**

To assess the impact of IFNα treatment, iEC were cultured in EGM-2 supplemented with IFNα (1000 U/mL) for 24 hours prior to harvesting, while negative controls were maintained in EGM-2 alone. Each condition was performed in triplicate for each genotype, resulting in a total of 18 samples generated across four independent experimental batches (Supplemental Table 7). Total RNA from purified iEC was isolated as described in the *RNA Extraction and Quantitative RT-PCR* section. All samples exhibited high RNA integrity, with RNA Integrity Numbers (RIN) ≥ 9, as determined using the TapeStation 4200 system (Agilent Technologies, Santa Clara, USA).

RNA concentration was measured using the Fluorometer Quantus™ (Promega, Fitchburg, USA). Following the manufacturer's protocol, ribosomal RNA was depleted using the NeBNext® rRNA Depletion Kit before library preparation with the NeBNext®Ultra™II Directional RNA Library Prep Kit for Illumina, both from New England BioLabs (Ipswich, USA), using an RNA input of 840 ng per sample. Subsequently, samples were sequenced in paired end reads (2x76 bp, dual indexed) on two NextSeq High Output Kits v2.5 (150 cycles) using a NextSeq 500 instrument from Illumina (San Diego, USA) to ensure sufficient reads for data analysis.

**Global RNA-sequencing Raw Data Processing**

Raw sequencing data in FASTQ format were generated using bcl2fastq (Illumina, San Diego, CA) for demultiplexing and conversion of base call (BCL) files. Subsequent RNA sequencing data processing was performed using the nf-core RNA-seq pipeline (v3.12)[6] with standard settings optimized for total RNA-Seq with ribosomal RNA depletion.

The pipeline included TrimGalore (v0.6.7)[7] for adapter trimming and quality filtering of reads. High-quality reads were then aligned to the reference genome (GRCh39) using STAR (v2.7.9a)[8] aligner under default parameters. Read counts at both gene and transcript levels were quantified using Salmon (v1.10.1)[9], allowing for accurate quantification of transcript abundance and accounting for biases such as GC content.

**Global RNA-sequencing Differential Expression Analysis**

The expression data were adjusted for unwanted technical and biological batch effects using the ComBat procedure. Differential expression analysis was conducted using the DESeq2 (v1.40.2) package in R (v4.3.0)[10]. Raw read counts were normalized, and differential gene expression between experimental conditions was determined. Genes with an adjusted p-value (FDR) < 0.01 were considered significantly differentially expressed.

To visualize the results, a Volcano plot was generated using the ggplot2 (v3.4.2) package in R[11]. The plot displays log2 fold changes versus the -log10 adjusted p-values, highlighting significantly upregulated and downregulated genes.

**Apoptosis Assay**

iEC were cultured in fully supplemented EGM-2 medium until confluency. To assess the effect of IFNα on apoptosis induction, IFNα (1000 U/mL) was added 48 hours prior to harvesting. A half-medium change was performed 24 hours after treatment initiation. Following enzymatic dissociation with trypsin, apoptosis was evaluated by flow cytometry using the FITC Annexin V Apoptosis Detection Kit with 7-AAD (BioLegend), according to the manufacturer's instructions.

**iEC Processing for Transmission Electron Microscopy**

iEC were cultivated in EGM-2 until confluency. After fixation with 3% glutaraldehyde for 1 hour at room temperature, iEC were dissociated using cell scrapers and processed for transmission electron microscopy (TEM). Following iPSC clones were used:

| Patient | Clone name | Genotype | Generated by |
|---|---|---|---|
| Patient 1 | PV1 007 WT | $JAK2^{WT}$ | Reprogramming |
| Patient 1 | PV1 009 HET | $JAK2V617F^{HET}$ | Reprogramming |
| Patient 1 | PV1 012 HOM | $JAK2V617F^{HOM}$ | CRISPR/Cas9 |
| Patient 1 | PV1 033 HOM | $JAK2V617F^{HOM}$ | CRISPR/Cas9 |
| Patient 2 | PV2 092 WT | $JAK2^{WT}$ | CRISPR/Cas9 |
| Patient 2 | PV2 070 HET | $JAK2V617F^{HET}$ | CRISPR/Cas9 |
| Patient 2 | PV2 021 HOM | $JAK2V617F^{HOM}$ | Reprogramming |

iEC were embedded in 5% low melting agarose (Merck) before being gelatinated and washed in 0.1 M Soerensen's phosphate buffer (Merck). They were then post-fixed in 25 mM sucrose buffer (Merck) containing 1% $OsO_4$ (Roth). Specimens underwent dehydration through an ascending ethanol series, with the final step repeated three times (30%, 50%, 70%, 90%, and 100% ethanol; 10 minutes each step). Following dehydration, samples were consecutively incubated in propylene oxide (Serva) for 30 minutes, in a mixture of EPON resin (Serva) and propylene oxide (1:1) for 1 hour, and in pure EPON for 1 hour. Polymerization of EPON was carried out at 90°C for 2 hours. Ultrathin sections of 90-100 nm thickness were produced using a UC6 ultramicrotome (Leica) equipped with a diamond knife (Diatome) and placed onto copper−rhodium grids (Plano). Samples were then viewed at an acceleration voltage of 60kV using a Zeiss Leo 906 (Carl Zeiss) transmission electron microscope.

TEM images from different genotypes were reviewed to assess potential ultrastructural differences. A trained person (Eva M. Buhl) from the Institute of Pathology and Electron Microscopy Facility (RWTH University of Aachen, Aachen, Germany) conducted the analysis in a blinded manner, ensuring that the reviewer was unaware of the genotype associated with each image. For each clone, one sample was processed for TEM, and per clone, 5 to 7 images were reviewed in a blinded manner. The assessment focused on key features such as the quantity of endoplasmic reticulum (ER), ribosomes, and Golgi apparatus, as well as structural clearness, which refers to the clarity and integrity of cellular ultrastructure, including the discernibility of organelle boundaries, membrane continuity, and the absence of artifacts. This approach ensured a robust identification of significant differences related to the genotypes under investigation.

**In Vitro Induction of Endothelial-to-Mesenchymal Transition**

Two-dimensional modeling of endothelial-to-mesenchymal transition (EndMT) was performed using JAK2$^{WT}$ and JAK2V617F iEC (passages 2–3). Cells were cultured in EGM-2 medium supplemented with 8 µM TGF-β inhibitor (StemCell Technologies) until reaching sub-confluency. iEC were then pre-incubated for 4 hours in starvation medium (EBM-2 supplemented with 2% FCS) containing one of the following conditions: IFNα (1000 U/mL), ruxolitinib (250 nM), nintedanib (250 nM), or a combination of ruxolitinib and nintedanib (both at 250 nM). EndMT was subsequently induced by the addition of a cytokine cocktail consisting of TGF-β (0.5 ng/mL), IL-1β (5 ng/mL), and TNF-α (10 ng/mL), as previously described[12]. After 48 hours of treatment, cells were enzymatically dissociated using trypsin, snap-frozen, and stored at -80 °C until RNA extraction.

**Generation of 3D Fibrin Hydrogels**

3D fibrin hydrogels were generated as previously described[13,14]. Following enzymatic dissociation, mesenchymal stromal cells (MSC) isolated from femoral heads of healthy donors, iEC, and iHC were resuspended in serum-free medium at the desired ratios (MSC:iEC = 3:1; MSC:iHC = 1:2). Cells were then mixed with aprotinin and thrombin, followed by the addition of fibrinogen. Detailed hydrogel compositions are provided in Supplemental Table 2.

Depending on the experimental setup, the resulting cell–matrix mixture was pipetted either into custom-fabricated cell crowns (produced in-house) placed within a 24-well plate format or directly into 8-well chamber slides (µ-Slide 8 Well high ibiTreat, Ref. 80806, Ibidi). Constructs were incubated to allow for polymerization and subsequently cultured under submerged conditions at 37 °C, 5% $CO_2$, and 100% humidity. Medium was partially replaced every 48 hours.

**Evaluation of the Anti-Angiogenic Potential of IFNα**

To visualize vascular tree formation within 3D fibrin hydrogels, MSC and iEC were fluorescently labeled using the PKH67 (green) and PKH26 (red) Cell Linker Kits (Millipore), respectively, according to the manufacturer's instructions. Labeled cells were subsequently embedded into 3D fibrin hydrogels and seeded into 8-well chamber slides (µ-Slide 8 Well, Ibidi), followed by culture in EGM-2 medium for up to 5 days.

To assess the anti-angiogenic effects of IFNα, two experimental approaches were employed. In the first, mono- and co-cultures were treated with IFNα (1,000 U/mL) on days 1 and 3 post-seeding. In the second approach, iECs and MSCs were pre-treated with IFNα (1,000 U/mL) prior to harvesting and co-embedding in 3D fibrin hydrogels. Within this setting, subsets of cocultures were either exposed to IFNα exclusively during the pre-treatment phase or received additional IFNα treatment on days 1 and 3 following hydrogel embedding.

Following fixation in 4% paraformaldehyde (PFA) overnight at 4 °C, nuclear staining was performed using 4',6-diamidino-2-phenylindole (DAPI; BioLegend). Slides were mounted with Dako Fluorescence Mounting Medium (Dako), and imaging was conducted using the automated EVOS M7000 Imaging System (Thermo Fisher Scientific). For each well, 25% of the total area was imaged, with identical positions selected across conditions to minimize investigator bias. Image analysis was performed using a custom macro developed in ImageJ to quantify the PKH26-positive fluorescence area. Representative images of spatial interactions between iEC and

MSC within 3D fibrin hydrogels were acquired using a Zeiss LSM 710 confocal microscope (Zeiss, Oberkochen, Germany). Image acquisition was performed using ZEN Black software, version 2.3 SP1 FP3 (64-bit).

**Modeling EndMT in 3D Assembloids**

EndMT modeling was performed using JAK2$^{WT}$ and JAK2V617F$^{HET}$ iEC (passage 2–3) expanded in EGM-2 medium supplemented with 8 µM TGF-β inhibitor. To enable tracking of vascular network formation during 3D coculture, iEC were pre-labeled with CellTracker CM-DiI (2 µM; ThermoFisher) prior to resuspension with MSC (passage 2–3) in fibrin hydrogels. Constructs were cultured in EGM-2 for 48 h. As in the 2D EndMT assay, 3D cocultures were pre-incubated for 4 h in starvation medium (EBM-2 + 2% FCS) supplemented with 1000 U/ml IFNα, 250 nM ruxolitinib, 250 nM nintedanib, or a combination of both inhibitors (250 nM each). EndMT was induced by addition of a defined pro-inflammatory and pro-fibrotic cytokine cocktail, as described above. Controls were maintained in starvation medium alone.
After 48 h, constructs were fixed in 4% PFA overnight at 4 °C and processed for paraffin embedding. Immunofluorescence staining was performed to assess expression of key EndMT markers (Supplemental Table 3).

**Generation of 3D Assembloids for Single-cell RNA Sequencing**

For single-cell RNA sequencing (scRNA-seq), 3D assembloids were generated by combining iMK, iM, and iGr (collectively referred to as iHC) at a 2:1:1 ratio. MSC ($6 \times 10^4$/ clot), iEC ($2 \times 10^4$/ clot), and iHC ($1.2 \times 10^5$/ clot) were harvested, resuspended in Iscove's Modified Dulbecco's Medium (IMDM; Thermo Fisher Scientific), and embedded in fibrin hydrogels as previously described.

On day 3, constructs were cultured in serum-free medium (SFM) supplemented with 2% FCS, SCF (50 ng/ml), TPO (10 ng/ml), VEGF (5 ng/ml), H-transferrin (6 µg/ml; Sigma), L-ascorbic acid (50 µg/ml; StemCell Technologies), and 1% penicillin/streptomycin. IFNα (1000 U/ml) was added for the final 48 h. Medium was refreshed every 48h. On day 5, fibrin hydrogels were enzymatically digested using a nattokinase-based protocol as previously described[15]. Briefly, nattokinase (HY-P2373; Hycultec) was reconstituted in 1 mM EDTA (in PBS) to a final concentration of 100 fibrin-degrading units (FU)/ml. Up to three hydrogels were incubated in 0.2 ml nattokinase solution at 37 °C for 30–45 min until fully dissolved.

**scRNA-seq of 3D Assembloids**

For the scRNA-seq experiment the 10x Genomics system was used and the manufacturers' instructions 'Chromium Next GEM Single Cell 3' Reagent Kitsv3.1 User Guide Rev D' were followed. For the present study, cells from 3D iEC-iHC-MSC cocultures were utilized. To enhance cell collection for scRNA-seq analysis, cells from three replicates per condition were pooled during the nattokinase digestion protocol. The following conditions were explored:

| | MSC | iEC | iHC | IFNα treatment |
|---|---|---|---|---|
| Sample 1 | $JAK2^{WT}$ | $JAK2^{WT}$ | $JAK2^{WT}$ | / |
| Sample 2 | $JAK2^{WT}$ | $JAK2V617F^{HET}$ | $JAK2^{WT}$ | / |
| Sample 3 | $JAK2^{WT}$ | $JAK2V617F^{HET}$ | $JAK2V617F^{HET}$ | / |
| Sample 4 | $JAK2^{WT}$ | $JAK2V617F^{HET}$ | $JAK2V617F^{HET}$ | 1000U/ml |

Following cell retrieval via nattokinase digestion and a PBS wash, samples were centrifuged at 500 x g for 10 minutes, and the supernatant was discarded. After resuspension in 1 ml PBS, cells were counted using trypan blue and a hemocytometer, with suspensions kept on ice throughout. The required cell suspension volume was calculated using a table from 10x Genomics, aiming for 8,000 cells per sample.

To enable transcriptomic analysis of single cells, Gel-Beads-in-Emulsion (GEMs) containing a Gel Bead (which barcodes each cell), a master mix (MM) for RT-PCR, and a single cell, were generated using the Chromium Controller (10x Genomics). After GEM generation, the samples were then transferred to a cycler ensuring dissolution of Gel Beads thereby releasing primers, and lysis of cells. The released primers contained an Illumina TruSeq Read1, 16nt 10xBarcode, 12nt unique molecular identifier and 30nt poly(dt)sequence. The incubation in the cycler then produced the full-length cDNA containing the barcoding of the individual cells.

After incubation in the cycler, 125µl Recovery Agent (10x Genomics) were added to each sample. Following discarding of the recovery agent and partioning oil, cDNA was purified using silane magnetic beads. After incubation with Dynabeads cleanup, tubes were placed on a magnet (10x Genomics) and the beads with the bound cDNA were held back, while the supernatant was discarded. After two washing steps with 80 % ethanol, magnetic beads were air dried, and cDNA was eluted using a freshly prepared elution solution according to manufacturer´s protocol and subsequently placed again on a magnet. Now the beads were retained, while cDNA was in solution, enabling the harvest of 35µl purified cDNA. In a next step the purified cDNA was further amplified by mixing it with 65µl of cDNA Amplification Reaction Mix and consecutive incubation in a thermal cycler as recommended in manufacturer´s instructions. After amplification the cDNA was again purified using magnetic beads, in this case the SPRIselect reagent (Beckman Coulter). After supernatant removal, beads were washed twice with 80 % ethanol while remaining on the magnet. The beads were air dried to remove remaining ethanol and the tubes were removed from the magnet. The beads were then suspended in EB buffer and incubated at room temperature to elute the cDNA. By placing the tubes on the magnet again, the beads were again retained, while 40µl purified cDNA were harvested in the supernatant.

**scRNA-seq Library Preparation and Sequencing**

Amplified cDNA was enzymatically fragmented to generate appropriately sized DNA fragments for Illumina sequencing. Adaptor ligation and sample indexing were performed according to the manufacturer’s instructions to enable multiplexing. Sequencing was carried out using the NovaSeq 6000 SP Reagent Kit v1.5 (100 cycles; Illumina, San Diego, USA) by the Genomics Facility of the Interdisciplinary Center for Clinical Research (IZKF), Faculty of Medicine, RWTH Aachen University.

**scRNA-seq Bioinformatical Analysis**

Bioinformatical analysis of the scRNA-seq data was performed by the IZKF genomics facility.

Prepared libraries were sequenced using the NovaSeq 6000 SP Reagent Kit v1.5. Raw outputs were demultiplexed using Cellranger mkfastq (v7.1, 10x Genomics). Reads were processed using the scRNA-seq nf-core pipeline (v2.5.1)[6]. Further data processing was performed using the Seurat package (v5.0)[16]. Low-quality cells were excluded by determining the percentage of mitochondrial genes for each cell and filtering out cells outside of 3 median absolute deviations from the median number of detected genes, read count, or mitochondrial genome content. Raw counts were normalized using the SCTransformV2 method from Seurat, and the effects of mitochondrial gene content and cell cycle differences were regressed out. The most variable 10% of genes were identified for downstream analysis.

Samples 1-4 were well mixed and did not require additional integration or harmonization after PCA and UMAP dimensional reductions. Cell clusters were detected using a modified Louvain algorithm for community detection on the neighborhood graph with a clustering resolution of 0.8. Individual clusters were manually annotated based on upregulated genes versus other clusters and the identification of canonical markers. Differential gene expression between detected clusters and between different samples within the same cluster was conducted using the MAST package wrapper in Seurat[17]. Genes with a log2 fold change $> 0.2$ and an adjusted p-value $< 0.05$ were considered significant. Additionally, differentially expressed genes were filtered to be expressed in at least 10% of the cells in either comparison group.

Overrepresentation analysis (ORA) and gene set enrichment analysis (GSEA) were performed using ClusterProfiler (v4)[18] and the Gene Ontology (GO) dataset. For pathway activity inference, we used the PROGENy[19] statistical model with the decoupleR package[20]. Each cell was scored for the activity of supported pathways using the top 500 genes from the PROGENy model. To compare pathway activity between samples, significant changes in score were evaluated using a t-test, and fold changes were computed from the generated scores. To compute a robust patch-independent enrichment score for individual gene sets, we used the AUCell package, which calculates the "Area Under the Curve" for the query gene set in the top 5% of expressed genes in each cell[21]. We calculated AUCell scores for the following signatures: Type I Interferon, JAK-STAT signaling pathway (KEGG), Hallmarks of the TGF-b Signaling (MsigDB: M5896), and extracellular matrix organization (GO:0030198). Additionally, a curated list of endothelial to mesenchymal transition genes was compiled from the literature.

To closely examine the stromal cell population, we independently identified and re-clustered CD45- or CD45low cells using single-cell expression of the PTPRC gene. This included endothelial cells, hemogenic endothelium, and pericytes. The stromal cells were clustered into 5 clusters. Differential gene expression and signature scoring were carried out as previously described. Differential composition analysis of endothelial cells was conducted using a permutation test with bootstrapping from the scProportionTest package[22]. All plots were generated using ggplot2[23] and ComplexHeatmap packages[24]. Unless otherwise described, statistical comparisons were conducted using the Student's t-test with Benjamini-Hochberg procedure for p-value correction.

**PV and MF Murine Models**

Experiments were conducted as part of a parallel project investigating IFNα effects on the BM niche *in vivo*. The JAK2V617F-driven PV murine model was previously established[25]. Briefly, FF1 JAK2V617F donor mice were Tamoxifen-induced (2 mg for 4 days) followed by 5 weeks phenotype development. Donor FF1 JAK2V617F and JAK2 WT BM mononuclear cells (BMMNCs) were transplanted in a 1:1 ratio into C57BL/6 recipient mice previously conditioned with radiation (2 x 4.5 Gy). Eight weeks after transplantation, mice were treated with 25 µg/Kg body weight pegylated IFNa (pIFNa) twice weekly (intraperitoneal administration) for up to 8 weeks. Blood parameters were monitored with the Element HT5 hematology analyzer (Heska) and at end point analysis, BM and spleen flow cytometry and BM scRNA-seq analysis was performed.

To model BM fibrosis, transplantation of thrombopoietin (ThPO)-overexpressing BM-derived cKIT+ hematopoietic stem and progenitor cells (HSPC) was performed as previously described[26]. Briefly, CD45.1 C57BL/6 BMMNCs were isolated and enriched for cKIT by MACS (Miltenyi Biotec) following manufacturer´s instructions. cKIT+ cells were cultivated in StemSpan supplemented with 50 ng/ml mSCF and 50 ng/ml mThPO (both Immunotools). Within 24h, ThPO-overexpression construct was delivered with lentivirus particles followed by additional 24h cultivation. Infected cells ($0.5x10^6$) were then washed and transplanted into irradiated (9 Gy) CD45.2 C57BL/6 recipient mice. Next, 28 days after transplantation, mice were treated with 25 µg/Kg body weight pIFNa twice weekly (intraperitoneal administration) for up to 6 weeks. Blood parameters were monitored with the Element HT5 hematology analyzer (Heska) and at end point analysis, BM and spleen flow cytometry and BM scRNA-seq analysis was performed.

**scRNA-seq of the JAK2V617F-driven PV murine model**

For the scRNA-seq experiment the 10X Genomics system was used and the manufacturers' instructions 'Chromium Next GEM Single Cell 3' Reagent Kitsv3.1 User Guide Rev D' were followed.

The scRNA-seq libraries were processed with the CellRanger pipeline (8.0.0) using the default parameters (transcriptome GRCm39-2024-A). Data integration and clustering were performed using Seurat (version 4.2.3). Cells with 400 or fewer expressed genes were removed, as were cells with more than 20,000 reads in total or more than 10% mitochondrial reads. Genes expressed in less than 5 cells were removed. Doublet detection and removal were performed using DoubletFinder[27]. Following data integration and principal component (PC) analysis, a shared nearest-neighbor graph was constructed using the first 50 PCs and clusters were determined with a resolution of 0.5 using harmony integration[28]. Cell types were annotated using published marker gene expression. Differentially expressed genes were determined using a MAST model[17].Only genes expressed in at least 10% of the cells were considered for comparison. Gene set enrichment analysis was performed using a univariate linear model as implemented by decoupler[20].

**scRNA-seq of the TPO-overexpression murine model of MF**

For the scRNA-seq experiment the 10x Genomics system was used and the manufacturers' instructions 'Chromium Next GEM Single Cell 3' Reagent Kitsv3.1 User Guide Rev D' were followed.

The scRNA-seq libraries were processed with the CellRanger pipeline (cellranger-7.1.0) using the default parameters (transcriptome mm10-2020-A). Data integration and clustering were performed using Seurat (version 4.2.3). Cells with 400 or fewer expressed genes were removed, as were cells with more than 20,000 reads in total or more than 20\% mitochondrial reads. Genes expressed in less than 5 cells were removed. Doublet detection and removal were performed using DoubletFinder[27]. Following data integration and principal component (PC) analysis, a shared nearest-neighbor graph was constructed using the first 50 PCs and clusters were determined with a resolution of 0.5 using harmony integration[28]. Cell types were annotated using published marker gene expression. Differentially expressed genes were determined using a MAST model[17]. Only genes expressed in at least 10% of the cells were considered for comparison. Gene set enrichment analysis was performed using a univariate linear model as implemented by decoupler[20].

**Bone Marrow Samples of IFNα-treated MPN Patients**

To validate IFNα´s anti-EndMT effects *in vivo*, BM trephines from MPN patients were analyzed prior to and during IFNα treatment. Patient characteristics are summarized in the table below. Abbreviations: ET, Essential Thrombocythemia; PV, Polycythemia Vera; HMR, High Molecular Risk; BMT, Bone Marrow Trephine.

| MPN Entity | Driver Mutation | HMR Mutation | Sex | Age | BMT 1st Timepoint | BMT 2$^{nd}$ Timepoint |
|---|---|---|---|---|---|---|
| ET | CALR | / | M | 60 | treatment-naive | IFNα treatment for approximately 12 months |
| PV | JAK2V617F | / | W | 38 | treatment-naive | IFNα treatment for approximately 5 months |
| ET | Triple negative | / | W | 64 | hydroxyurea | IFNα treatment for approximately 24 months |

**Histology and Immunofluorescence Staining**

Hematoxylin and eosin staining was performed to assess the structural organization of 3D assembloids. Immunofluorescence staining was used to evaluate protein expression of selected markers (see Supplemental Table 3). Paraffin sections were deparaffinized, rehydrated in graded ethanol, subjected to heat-mediated antigen retrieval in citrate buffer (Vector Laboratories), and blocked with 5% goat serum in PBS. Primary antibodies were incubated overnight at 4 °C. Duplex IF staining for FSP1 and CD31 was performed to assess colocalization. The following day, secondary antibodies were applied for 1 h at room temperature, followed by nuclear counterstaining with DAPI. Slides were mounted using Dako Fluorescence Mounting Medium.

**Image Acquisition and Quantification**

Fluorescence imaging was performed using the EVOS M7000 system. Mean gray value or fluorescent area per regions of interests (ROI) was quantified using ImageJ. To minimize investigator bias, three predefined regions of interest (ROIs; 1392 × 1320 pixels) were selected per condition based on nuclei density in the DAPI channel. For analysis of IF staining in human and murine BM samples, vessel structures within the medullary space were identified by nuclear morphology (DAPI) and CD31 positivity and outlined as ROIs. FSP1 mean gray value was measured and normalized to DAPI counts. Colocalization of FSP1 and CD31 signals was assessed with the Just Another Co-localization Plugin (JaCoP) for ImageJ[29].

**Supplemental Figure 1. Exploring neoplastic vasculogenesis using MPN patient-derived iEC**

(A) The purity of isolated CD144+ cells was assessed by FACS for endothelial markers (CD144, CD31, CD105) and the hematopoietic marker CD45. One-way ANOVA (n = 3).

(B) For proliferation assays, iEC were seeded at a density of 20,000 cells per well in a 24-well plate, and cell counts were determined after 4 days of culture in supplemented Endothelial Growth Medium-2. One-way ANOVA (n = 5).

(C) To compare the angiogenic potential of mutated iEC to WT control, ETFA were performed. Images were taken 12 to 14 hours after seeding the cells onto Matrigel® or Geltrex® in a 96-well plate format. Scale bars: 200 μm.

(D) ETFA quantitative analysis comparing the total length of tubule-like structures formed across three genotypes under basal conditions. One-way ANOVA (n = 3).

(E) To assess the response of iEC to clinically used or compounds under investigation for MPN therapy, iEC were treated with 1 μM of the listed compounds or 1000 U/ml IFNα. One-way ANOVA (n = 3-7).

(F) Experimental design used to explore the anti-angiogenic potential of IFNα. ETFA were conducted with or without prior exposure of iEC to 1000 U/ml IFNα before seeding onto reconstituted basement membrane.

(G) Quantitative analysis of the impact of IFNα treatment on the total length of capillary-like structures formed by iEC. Statistical significance was assessed using a paired t-test (n = 3-5).

(H) Photographs illustrating the effects of 1000 U/ml IFNα pre-sensitization on iEC. White arrows highlight disrupted tubule-like structures. Scale bars: 100 μm.

(I) Quantitative analysis of ETFA comparing the total length of capillary-like structures following iEC pre-treatment with 1000 U/ml IFNα. Unpaired t-test (n = 3).

(J) Quantification of the impact of IFNα treatment on IFNα pre-treated iEC. Paired t-test (n = 3).

**Supplemental Figure 2. IFNα elicits a core interferon response without triggering apoptosis in iEC**

(A) iEC demonstrate high cosine similarity values with primary human EC. A one-sided Wilcoxon test was employed to determine whether the cosine similarity values for iEC were significantly greater than those of other cell types.

(B) Normalized read counts of mRNA expression levels for JAK2 and STAT1 from global RNA-seq in JAK2V617F and JAK2$^{WT}$ iEC. One-way ANOVA (n = 3).

(C) Gene Ontology (GO) analysis highlighting iEC core IFNα response. The size of the dots reflects the number of dysregulated genes from the gene list associated with the GO term, while the color of the dots represents the adjusted p-value (p. adjust).

(D) Quantification of Interferon-Induced Protein with Tetratricopeptide Repeats 1 (*IFIT1*), Signal Transducer and Activator of Transcription 1 (*STAT1*) and Ubiquitin-Specific Peptidase 18 (*USP18*) mRNA levels by RT-qPCR in treated versus untreated iEC. Paired t-test (n = 6).

(E) Apoptosis induction following IFNα treatment was evaluated across the three genotypes using Annexin V/7-AAD staining and analyzed by FACS. Representative box plot illustrating the used gating strategy.

(F) Quantification of the percentage of Annexin-V+/ 7-AAD+ iEC among the three genotypes, following 48-hour treatment with IFNα. One-way ANOVA (n = 3).

(G) PROGENy analysis of JAK–STAT pathway activity in iEC following IFNα treatment.

**Supplemental Figure 3. JAK2V617F$^{HET}$ iEC exhibit a pro-mesenchymal phenotype while JAK2V617F$^{HOM}$ iEC display dysregulated translation machinery**

(A) Quantification of normalized read counts for mRNA expression levels of Small Nucleolar RNA, H/ACA Box 73B (*SNORA73B*), RNA, Y3 (*RNY3*), RNA, U11 small nuclear (*RNU11*), and RNA, U5, small nuclear 1 (*RNU5E-1*) in JAK2V617F$^{HOM}$ iEC compared to their WT controls (n = 3).

(B) GO analysis of terms significantly downregulated in JAK2V617F$^{HOM}$ versus JAK2$^{WT}$ iEC. Dot size indicates the number of dysregulated genes associated with the GO term, while dot color represents the adjusted p-value (p. adjust).

(C) TEM images illustrating reduced rough endoplasmic reticulum (rER) abundance in JAK2V617F$^{HOM}$ iEC. Features highlighted include cell nucleus (N), loose ribosomes (red arrow), and rER (white arrow). Scale bar: 1,000 nm.

(D) Quantification of normalized mRNA read counts for RNA, U12 small nuclear (*RNU12*) and Small Nucleolar RNA, H/ACA Box 23 (*SNORA23*) expression in untreated versus IFNα-treated JAK2$^{WT}$ (n = 3) and JAK2V617F$^{HOM}$ iEC (n = 8). Paired t-test.

(E) List of significantly dysregulated hallmark gene sets in JAK2V617F$^{HET}$ iEC (cut off: nominal p-value <0.05 and false discovery rate <0.25). NES, normalized enrichment score.

(F) GO analysis of genes significantly upregulated in JAK2V617F$^{HET}$ iEC compared to JAK2$^{WT}$ iEC. Dot size indicates the number of dysregulated genes associated with the GO term, while dot color represents the adjusted p-value (p. adjust).

(G) Quantification of normalized mRNA read counts for Bone Morphogenetic Protein 4 (*BMP4*) and Fibronectin 1 (*FN1*) expression in JAK2V617F$^{HET}$ iEC compared to JAK2$^{WT}$ (n = 3).

(H) Quantification of normalized mRNA read counts for *FSP1*, *ACTA2*, *BMP4* and *FN1* expression in untreated versus IFNα-treated JAK2V617F$^{HET}$ iEC (n = 3).

**Supplemental Figure 4. EndMT is suppressed by TKIs and IFNα in 3D iPSC-based assembloids**

(A) Schematic overview of the experimental workflow used to generate 3D assembloids for modeling neoplastic vasculogenesis. MPN patient-derived iEC and primary MSC were embedded in fibrin hydrogels as either monocultures or cocultures. Representative immunofluorescence image of a 3D assembloid stained for CD31 (green) to visualize iEC; nuclei are counterstained with DAPI (blue). An asterisk (*) indicates a lumen-like structure. Scale bar: 25 µm. Representative transmission electron microscopy image shows a vessel-like structure with annotated features: lumen (*), abluminal extracellular matrix (orange box), and primary cilium (red box). Scale bar: 5,000 nm. These 3D systems provide a scalable platform for disease modeling and *in vitro* drug screening.

(B) 3D culture medium was supplemented with 1000 U/mL IFNα on days 1 and 3. After 5 days of incubation, samples were fixed and analyzed by microscopy. Vascular network formation was assessed by quantifying the PKH26 fluorescence area in 3D iEC monocultures and iEC-MSC cocultures under different genetic and treatment conditions. Statistical analysis was performed using one-way ANOVA (n = 3-4).

(C) Experimental layout for investigating the impact of IFNα pre-treatment on vasculogenesis. Before 3D coculture, MSC and iEC were separately pre-treated with IFNα (1000 U/mL). Vascular network formation was assessed by quantifying PKH26 fluorescence, comparing the effects of separate iEC and MSC pre-treatment with 1000 U/mL IFNα on the total area of capillary-like structures. Statistical analysis was performed using an unpaired t-test (n = 3).

(D) Representative IF images of JAK2V617F$^{HET}$ assembloids under basal conditions and after pro-fibrotic cytokine exposure, stained for collagen1a1 (Col1a1, green) with DAPI (blue) highlighting cell nuclei. Quantitative analysis of mean fluorescence intensity (MFI). Two-way ANOVA (n = 3). Scale bars: 250 µm.

(E) Quantification of mRNA levels of Snail family transcriptional repressor 1 (*SNAI1*), Platelet endothelial cell adhesion molecule 1 (*PECAM1*), and Claudin 5 (*CLDN5*) by RT-qPCR in cytokine-treated JAK2$^{WT}$ and JAK2V617F$^{HET}$ iEC cultured in 2D monolayers. Two-way ANOVA (n = 4).

(F) CD31 fluorescence area quantification in cytokine-treated 3D assembloids pre-treated with IFNα, nintedanib, ruxolitinib, or the combination of ruxolitinib and nintedanib. One-way ANOVA (n = 3).

**Supplemental Figure 5. scRNA-seq of a 3D iPSC-derived BM niche model**

(A) Characterization of iPSC-derived hematopoietic stem or progenitor cells (iHSPC; CD117+CD34+), granulocytes (iGr; CD45+CD66b+), megakaryocytes (iMK; CD45+CD61/41+), and monocytes/macrophages (iM; CD45+CD68+) by cytospins and FACS. Scale bars: 10 µm.

(B) UMAP plot showing the subclustering of non-hematopoietic ($CD45^{-}$) lineages and (b) their distribution across cell cycle phases (G1, S, and G2). Abbreviations: MSC, mesenchymal stromal cells; EC, endothelial cells; HemoEndo, hemogenic endothelium; Peri, pericytes.

(C) Circle plot illustrating the top 25% of ligand-receptor (L-R) interaction strength among various cell types in $JAK2^{WT}$ assembloids. Each node represents a distinct cell type, with MSC clusters classified as stromal cells. Line thickness reflects interaction strength, and arrows indicate the direction of interactions.

(D) Flow charts illustrating the information flow derived from L-R interaction analysis in $JAK2^{WT}$ assembloids, specifically focusing on endothelial cells, stromal cells, and pericytes.

**Supplemental Figure 6. JAK2V617F hematopoietic cells promote niche pro-fibrotic reprogramming**

(A) Heatmap showing the dysregulation of EndMT-associated genes in $JAK2V617F^{HET}$ iEC. Data derived from global RNA-sequencing analysis.

(B) Lollipop plot showing expression of EndMT-associated genes in EC from $JAK2V617F^{HET}$ (HET-HET) compared to $JAK2^{WT}$ assembloids (WT-WT).

(C) Circle plot illustrating the overall L-R interaction strength among various cell types in 3D assembloids with vascular-restricted $JAK2V617F^{HET}$ (HET-WT) compared to $JAK2^{WT}$ assembloids (WT-WT). Each node represents a distinct cell type, with MSC clusters classified as stromal cells. Line thickness reflects interaction strength, and arrows indicate the direction of interactions. Red lines represent increased interaction strength, while blue lines indicate decreased interaction strength.

(D-E) Circle plots illustrating MIF–CD74/CD44 (D) and LGALS1–ITGB1 (E) interactions in JAK2V617F$^{HET}$ and JAK2$^{WT}$ assembloids. Nodes represent distinct cell types; MSC clusters are grouped as stromal cells. Line thickness reflects interaction strength; arrows indicate interaction direction.

**Supplemental Figure 7. Assembloids recapitulate MF-associated transcriptional reprogramming of stromal cells**

(A) Heatmap showing pathway enrichment across stromal cell subsets under different experimental conditions. Columns represent pathways; rows denote distinct stromal clusters.

(B-C) Box plots showing AUC scores for (B) TGFβ signaling and (C) ECM-related pathways in non-hematopoietic (CD45$^{-}$) populations from 3D assembloids. Color code indicates assemboids genotype and treatment: blue, WT iEC–WT iHC; yellow, HET iEC–WT iHC; orange, HET iEC–HET iHC; green, HET iEC–HET iHC treated with IFNα (1000 U/mL, 48 h).

**Supplemental Figure 8. Investigating the impact of pegylated IFNα on neoplastic vasculature in a JAK2V617F-driven PV mouse model**

(A) Schematic overview of the workflow for EC isolation from murine BM. Following mechanical dissociation and enzymatic digestion, cells were processed by FACS to isolate CD31$^{+}$CD45$^{-}$Ter119$^{-}$ EC for subsequent scRNA-seq.

(B) Peripheral blood counts of white blood cells (WBC), neutrophils, red blood cells (RBC), hemoglobin (HGB) levels, and percentage of Lin$^{-}$Sca1$^{+}$ progenitor cells in the BM of WT and JAK2V617F mice following 4 weeks of pegylated IFNα (pIFNα) treatment. Two-way ANOVA (n = 6).

(C) Representative reticulin stain images of BM trephines from untreated and 8-week pIFNα-treated WT and JAK2V617F mice. Scale bars: 150 μm.

(D-E) Heatmaps showing the expression of canonical endothelial markers (D) and vascular sprouting–associated genes (E) across EC clusters identified by scRNA-seq.

**Supplemental Figure 9. pIFNα-induced vascular niche reprogramming in a JAK2V617F PV Mouse Model**

(A) GSEA heatmap showing hallmark pathway enrichment across endothelial subtypes (arteriolar, arterial, BBB/CNS-like, and sinusoidal EC) from scRNA-seq of BM-derived EC 12 weeks post-engraftment. Comparisons: JAK2V617F vs. WT (–), and JAK2V617F + pIFNα vs. JAK2V617F (+).

(B) GSEA of cytoplasmic translation (C. T.) and ribosome-associated pathways (Rib.) in endothelial subclusters from JAK2V617F vs. WT mice following 8 weeks of vehicle or pIFNα treatment.

**Supplemental Figure 10. Transcriptional profiling of endothelial subtypes in a TPO-overexpression MF mouse model**

(A) Representative reticulin stain images of BM trephines from untreated and 4-week pIFNα-treated WT and TPO mice. Scale bars: 150 μm.

(B) UMAP of $CD31^{+}CD45^{-}Ter119^{-}$ EC from BM of the TPO MF murine model. Cells are clustered into 7 transcriptionally distinct endothelial populations based on marker gene expression, including 4 arterial subclusters: (1) vascular remodeling, (2) vessel sprouting, (3) $Snai2^{+}Twist2^{+}$, and (4) progenitor-like EC; and three non-arterial subsets: (5) arteriolar, (6) sinusoidal, and (7) BBB/CNS-like EC. Cell cycle phase (G1, S, G2M) are overlaid in the right and bottom panels.

(C) Heatmaps showing the expression of canonical endothelial markers and vascular sprouting–associated genes across EC clusters identified by scRNA-seq.

(D) Dot plot showing expression of selected genes across endothelial clusters from WT and TPO mice ± 4-week pIFNα treatment. Genes are grouped by functional annotation: endothelial markers (blue), and vascular remodeling/EndMT-associated genes (orange).
